# Cell-Based Immunization Combined with Single-Round Cell Panning Enables Discovery of PSMA-Targeting Nanobodies from Phage Display Libraries

**DOI:** 10.1101/2025.11.17.688677

**Authors:** Tong Yang, Joke Veldhoven-Zweistra, Maarten Ligtenberg, Sigrun Erkens, Mirella Vredenbregt-van den Berg, Rick Jansen, Patrick Chames, Eric M.J. Bindels, Khadijeh Ahmadi, Chris H. Bangma, Anton M. F. Kalsbeek, Janne Leivo, Nicolaas Lumen, Harmen J. G. van de Werken, Wytske M. van Weerden, Soudabeh Kavousipour, Raheleh Tooyserkani, Guido Jenster

## Abstract

There is a strong need for nanobodies targeting novel cancer-associated antigens to advance radioligand imaging and antibody-based therapeutics. In this study, we investigated whether non-targeted Llama immunization using tumor cells, combined with non-targeted phage display panning on human cell lines, could yield nanobodies specific to Prostate Specific Membrane Antigen (PSMA). Nanobody selection using both classical three round PSMA negative-positive panning and single round panning of cell lines positive or negative for PSMA showed clear enrichment for PSMA binders in both strategies. Using shRNA knockdown, flow cytometry, cell-ELISA, immunohistochemistry and structural modeling and docking, we confirmed the PSMA targeting of selected nanobodies. Two distinct epitopes were predicted to be bound by nanobodies PSMANb9 and A7 (JVZ-007), corroborated by epitope competition assays. These findings support the feasibility of non-targeted immunization and panning strategies for isolating clinically relevant cancer nanobodies.

## 1. Introduction

The 2018 Nobel prize winning phage display technology was first introduced by George Smith in 1985 for peptide display [1] and later exploited by McCafferty et al. to display antibody fragments [2]. The most commonly used antibody fragments include the single-chain variable fragments (scFvs) and VHHs (also known as nanobodies). ScFvs are engineered fusion proteins of the variable regions of both heavy and light chains, while VHHs are naturally occurring, single-domain antibody (sdAb) fragments derived from camelid heavy-chain antibodies [3]. Libraries of VHH antibodies are typically generated by immunization of Llamas with the antigen of interest, after which multiple rounds of pannings take place against the immobilized single target protein [4]. Next Generation Sequencing (NGS) of > 10^5^ selected sdAbs replaced the classical Sanger sequencing of <1000 individually picked clones, which had a major impact on insights into enrichment and binding characteristics of clonally related sequences [5].

In our search for nanobodies against cancer, we questioned whether non-targeted Llama immunization using tumor cells and non-targeted panning using cell lines could identify nanobodies against a specific cell surface antigen. As our target of interest, we selected Prostate Specific Membrane Antigen (PSMA), a type II transmembrane glycoprotein, also known as glutamate carbopeptidase II (GPCII). PSMA is highly expressed in normal prostate epithelial cells and additionally in brain, small intestine, salivary glands and kidney [6]. In most primary prostate cancer (PCa) and particularly in advanced metastatic lesions, PSMA is overexpressed and is a well-known target for nuclear imaging [7–9] and radioligand therapeutics [10, 11]. Particularly PSMA PET (Positron Emission Tomography) tracers (e.g. 68Ga-PSMA-HBED-CC (PSMA-11), 68Ga-PSMA-617, 68Ga-PSMA-I&T) changed clinical diagnostics of metastasized PCa.

We and others previously identified nanobodies against PSMA and tested their efficacy in imaging as mono- and bivalent nanobodies and in therapeutics as bispecifics and CAR-T fusions [12–19]. The clinical application of nanobodies is strengthened by their unique structural features including their small size (~15 kDa), enhanced tissue penetration and high stability [20]. Technology-wise, the full nanobody sequence (110-140 amino acids) can be determined for millions of clones in a straight-forward NGS run. Having access to the increasing number of nanobody sequences and their target proteins, fuels research on VHH structural modelling and docking [21, 22]. The next steps towards AI-driven target prediction and *de novo* nanobody design are being taken and will strongly benefit from large panning databases [23, 24].

In this study, we compared panning protocols of nanobodies from 3 phage display libraries to identify PSMA binding nanobodies. A single panning round against a series of human cell lines was compared to classical 3-round negative-positive targeted panning. Candidate nanobodies exhibiting PSMA-associated binding were examined in cell binding assays and further validated in immunostaining of human normal and cancer tissues. Among these, specificity for PSMA of PSMANb9 and A7 (JVZ-007) [12] were confirmed in PSMA knockdown LNCaP cells. *In silico* analysis of nanobody-PSMA interactions revealed two distinct binding sites on the extracellular domain of PSMA and confirmed by *in vitro* epitope competition assays.

## 2. Results

### 2.1 Schematic workflow of the panning and selection strategy

In our search for an undirected cell-based panning approach to identify novel PSMA nanobodies, we compared two panning strategies: (i) the classical targeted panning against the wild type mouse melanoma B16 (B16-WT) cell line and human PSMA-expressing B16 (B16-PSMA) cell line and (ii) panning against a panel of PSMA-positive and PSMA-negative human cell lines (Suppl. Figure 1).

For the nanobody phage panning, we included three different libraries. L1P4 was previously generated by immunization of a Llama with the prostate cancer (PCa) cell line suspensions of (PSMA-positive) VCaP, PC346C, LNCaP and MDAPCa2b [12]. The LUPCa1 and LUPCa2 libraries were created by Llama immunization with cell fractions from a collection of fresh frozen PCa and bladder cancer (BlCa) patient tumors (Materials and Methods Table 1).

**Table 1.**
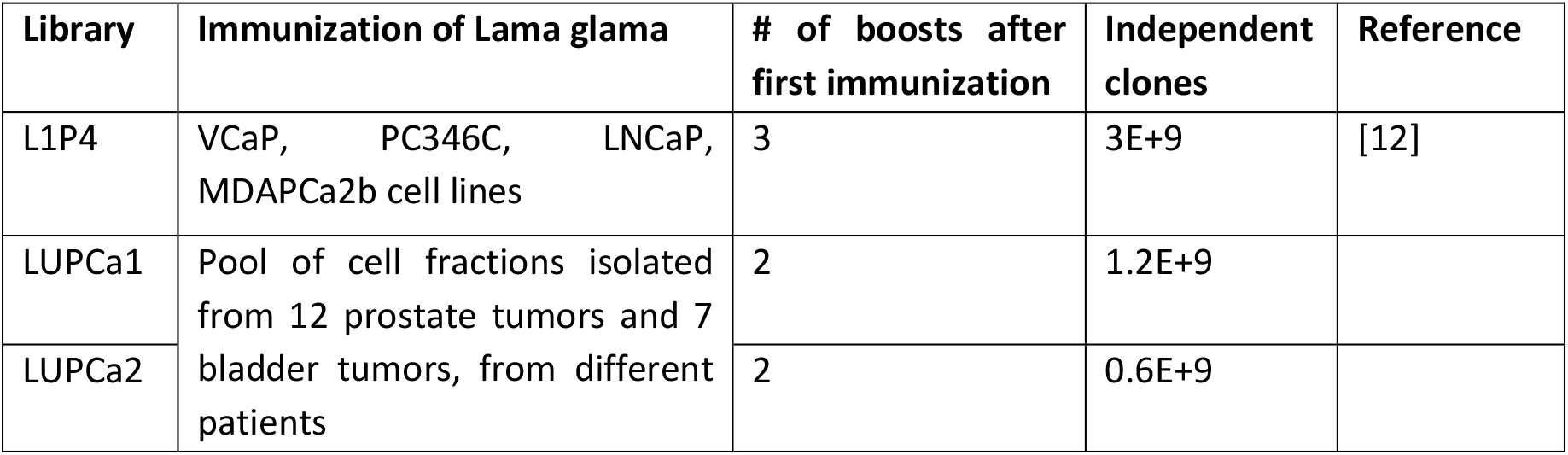
Immunizations and characteristics of the three nanobody-phage display libraries.

### 2.2 PSMA-targeted L1P4 and mixed library panning using B16-WT and B16-PSMA cell lines

For the targeted panning against B16-WT and B16-PSMA using the L1P4 library, we compared two different panning procedures: (i) classical 3 rounds of negative-positive panning (3R-NegPos) and (ii) 1 round single cell line panning (1R-SC).

The 3R-NegPos selection consisted of three rounds, starting with 100x L1P4 library against B16-WT cells, followed by incubation of the unbound fraction with B16-PSMA cells to isolate bound nanobody-phages. Upon next generation sequencing (NGS) and database building, we selected 462 nanobody clusters with different CDR3s that were >5-times more abundant in round 2 and 3 as compared to round 1, with a minimal read count of 0.001% in round 3 of the total number of reads.

To select PSMA-targeting nanobodies without negative selection, we performed a 1 round single cell line panning (1R-SC) using B16-WT and B16-PSMA cells. We selected 341 nanobody clusters that were >5-times more abundant in the B16-PSMA panning as compared to B16-WT and had a read count of >0.001% in B16-PSMA of the total number of reads.

The overlap between the 462 3R-NegPos nanobody clusters and the 341 1R-SC clusters was only 6 (p<1.7*10^−4^) (Figure 1A). These 6 common nanobody clusters were strongly enriched during 3R-NegPos panning from 0.001% of the reads in the L1P4 library to 3.0%, 15.7% and 29.9% of the reads in round 1, 2 and 3, respectively. In the single round B16-PSMA panning, the 6 PSMA nanobody clusters accounted for only 0.41% of the total number of reads sequenced. Previously, using 3R-NegPos panning, the A3, A5 (JVZ-005) and A7 (JVZ-007) PSMA nanobodies were identified via colony-picking [12]. This corresponds to our NGS identification as these 3 nanobodies overlap with the 6 highly abundant clusters. The other 3 abundant nanobody clusters are all highly similar to the A3 CDR3 sequence and are expected to bind the same PSMA epitope.

**Figure 1:**
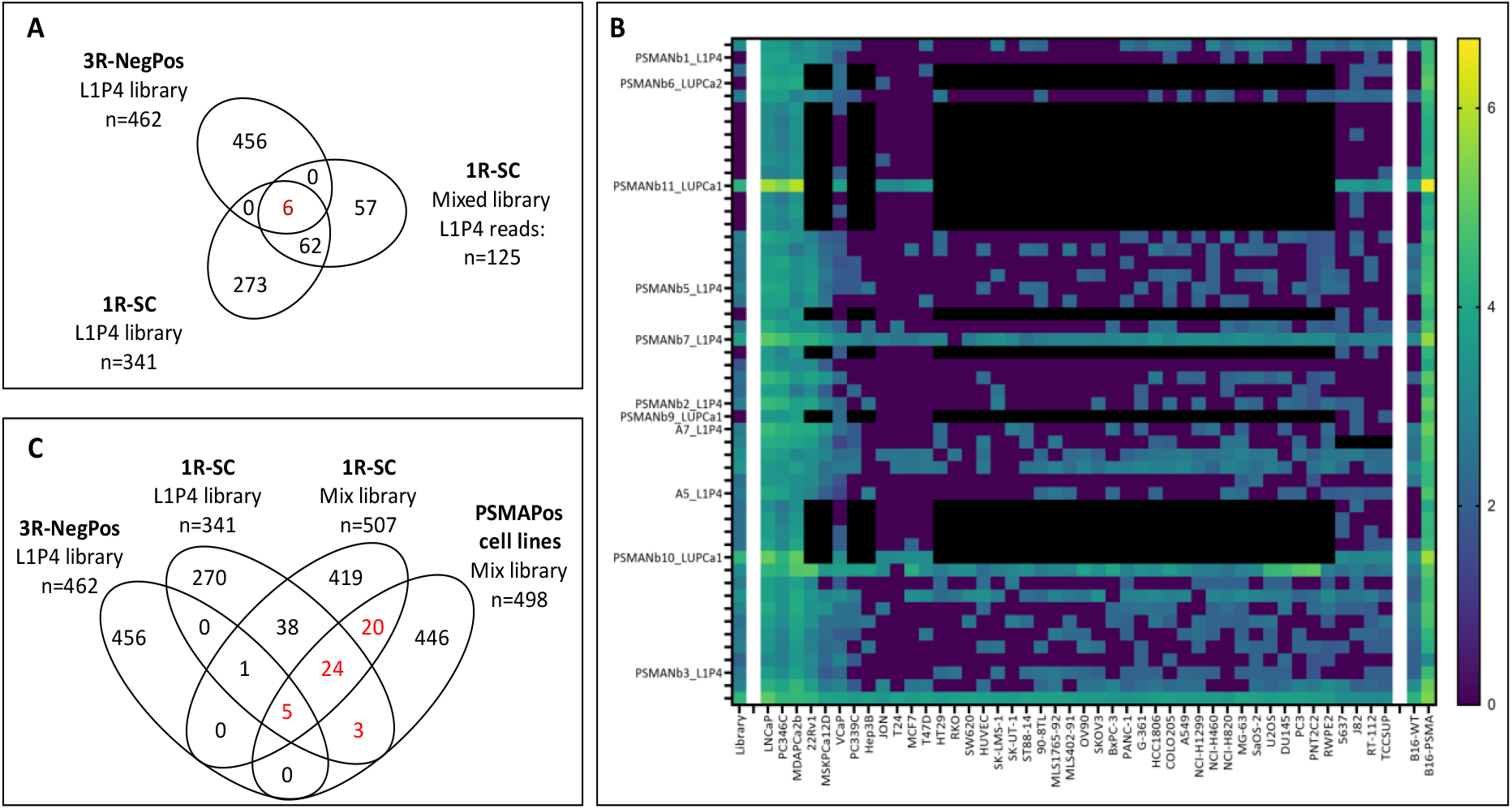
**(A)**: Venn diagrams of the overlap between panning rounds. Comparisons of the 3-round negative-positive (3R-NegPos) panning against B16-WT and B16-PSMA and the 1 round single cell (1R-SC) pannings using the L1P4 and mixed nanobody-phage libraries. **(B)** Heatmap of log10(n+1) values of 52 nanobody clusters preferentially binding to a series of PSMA-positive cell lines. The first column represents the abundance of the nanobody clusters in the original library followed by abundance after 1 round of panning against different (cancer) cell lines. Cell lines are ordered from left-to-right based on their FOLH1 RNA expression and binding of a PSMA antibody. Final two columns represent binding to B16-WT and B16-PSMA in a 1 round single cell panning. The 11 selected nanobodies for validation are depicted. Black: missing value. **(C)** Comparisons of the targeted pannings against B16-WT and B16-PSMA with the 1 round panning against a series of PSMA positive (PSMApos) and negative cell lines. The overlapping 52 nanobody clusters were selected for further validation (depicted in red).

Next to the panning using the L1P4 library, we tested whether a mix of L1P4 with the LUPCa1 and LUPCa2 libraries could reproduce the 1R-SC using the B16-WT and B16-PSMA pannings. The 3 libraries were mixed and based on a vector barcode, the origin of each individual nanobody upon sequencing was determined.

Using an identical 1R-SC panning protocol and nanobody selection criteria, 507 nanobody clusters were selected as higher binders in B16-PSMA cells as compared to B16-WT. Of these 507 nanobodies, 125 were from L1P4 of which 68 overlap with the 341 nanobody clusters identified using the L1P4 library mentioned above (p<1*10^−150^). All 6 nanobody clusters that overlapped with the 3R-NegPos were present in the list (Figure 1A).

### 2.3 Single round panning against a collection of human cell lines

Next, we performed 1 round of panning using the L1P4 library against 40 different (cancer) cell lines and the mix of 3 libraries (L1P4 with LUPCa1 and LUPCa2) against 12 cell lines. The cell lines were grouped in PSMA positive (PSMApos) and PSMA negative (PSMAneg) cell lines based on the level of FOLH1 mRNA (encoding the PSMA protein). We selected 498 nanobody clusters with on average more than 5-fold higher binding to PSMApos cell lines and a minimal read count of 0.001% of the average count of the PSMApos cells. Comparison with the 3R-NegPos and 1R-SC selections against B16-PSMA, identified an overlap of 52 nanobody clusters of which 5 intersect with all groups and 47 with 1R-SC (Figure 1B and C; Suppl. Table 1).

The combined targeted B16-PSMA and undirected cell line pannings resulted in the identification of 32, 18 and 2 PSMA-binding nanobody clusters from the L1P4, LUPCa1 and LUPCa2 libraries, respectively. From these groups, we selected 11 nanobody clusters (A5, A7, and PSMANb1-3, 5-7, 9-11) based on binding specificity to B16-PSMA and PSMApos cell lines, their panning procedure and library origin (Figure 1B). The most abundant nanobody sequence from each cluster was DNA synthesized, cloned into the pHEN vector and phages were regenerated in TG1 bacteria for nanobody-phage validation. Although the clone-picked A3 nanobody did not fulfil the PSMApos/PSMAneg cell line selection criteria (<0.001% of reads), we took it along as it was identified by colony-picking previously. In total 12 nanobody-phages were generated for further validation.

### 2.4 Circular dendrogram illustrating library-specific clustering of nanobody CDR3 repertoires

Circular dendrogram analysis was performed to visualize the sequence diversity and inter-library distribution of nanobody CDR3 repertoires after 1R-SC PSMApos cell line panning (n=498; Suppl. Figure 2A). In the mixed repertoire, distinct sequence families corresponding to the three nanobody libraries were evident. The L1P4 library displays the highest diversity, harboring numerous potential PSMA-targeting nanobodies with low sequence similarity. In contrast, the LUPCa1 library contains a large number of sequences mainly confined within several dominant families, whereas most of the 52 overlapping PSMANbs fell outside these clusters. The LUPCa2 library exhibited overall fewer PSMApos cell line nanobody sequences, reflecting inter-animal variability in immune response. In the refined dendrogram of the 52 overlapping clusters (Suppl. Figure 2B), nanobodies from all three libraries were represented, including both unique and family-associated members, collectively illustrating the repertoire’s sequence diversity and representativeness for subsequent experimental validation.

### 2.5 Validation of binding specificity of selected PSMA nanobodies

We tested the binding specificity of the selected 12 nanobody-phages in a whole cell-ELISA assay using 4 cell lines: PSMA-positive cell lines (LNCaP and B16-PSMA) and PSMA-negative cell lines (DU145 and B16-WT) (Figure 2A). Six out of twelve nanobodies (A3, A7, and PSMANb5, 6, 9, and 11) showed significant higher binding in PSMA positive versus negative cell lines. We further examined PSMA-binding specificity of these 6 nanobody phages against the IgG PSMA antibody. 20 different PSMApos and PSMAneg cell lines were used in an immunohistochemistry (IHC) assay where nanobody-phages were incubated with cells spotted on a glass slide (Figure 2B; Suppl. Figure 3).

**Figure 2:**
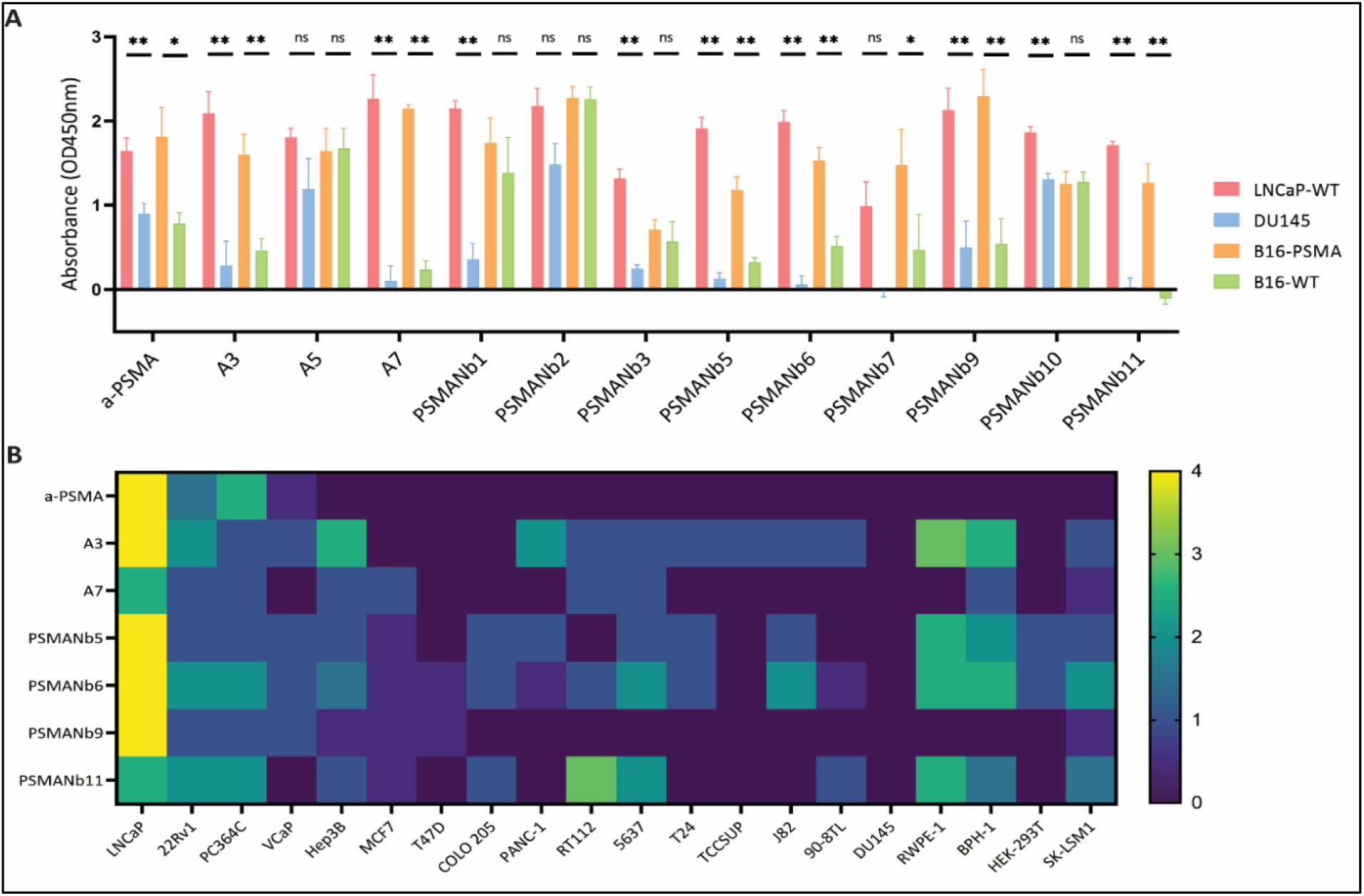
Validation of binding of selected nanobodies to cell lines in cell-ELISA and immunohistochemistry. **(A)** Nanobody-phage binding to PSMA negative (B16-WT, DU145) and PSMA positive (B16-PSMA, LNCaP) cell lines in a whole cell-ELISA assay. A regular anti-PSMA (a-PSMA) antibody was used as positive control. ELISAs were performed twice in triplicate. Absorbance values are represented as mean ± SEM values. Nonparametric Mann-Whitney test, *p<0.05, **p<0.01, ***p<0.001, ns: non-significant. **(B)** Heatmap representation of PSMA nanobodies and a PSMA antibody binding to spotted cells from 20 different cell lines using immunohistochemistry. Blinded scoring to a scale of 0-4 (Suppl. Figure 3) was performed independently by two scientists.

The validation reproduced the panning data well: Pearson Correlation Coefficient of the IHC scores and panning data of 6 nanobody-phages was on average (±SD) 0.60 (±0.18). As expected, the IgG anti-PSMA antibody bound specifically to PSMA positive cell lines with a binding level reflecting the known FOLH1 RNA expression levels. The only exception was the VCaP cells having high mRNA levels, but relatively low antibody surface binding. Among the nanobody-phages, A7 and PSMANb9 consistently yielded strong staining across PSMA-positive cell lines, with the most intense staining observed in LNCaP cells. As compared to A3, A5 and PSMANb5, 6, 11, the A7 and PSMANb9 nanobodies only bound (weakly) to a few PSMA negative cell lines.

### 2.6 Nanobody phages A7 and PSMANb9 specifically bind prostate (cancer) tissue

To further investigate the PSMA specificity of A7 and PSMANb9, we stained frozen sections of human PCa and normal adjacent prostate (NAP) tissues. IgG anti-PSMA antibody strongly stained PCa tissues compared to NAP tissues (Figure 3A, E). Similarly, A7 and PSMANb9 nanobody-phages exhibited comparable stronger staining patterns in PCa (Figure 3B, C) versus NAP, although with weaker intensity (Figure 3F, G).

**Figure 3:**
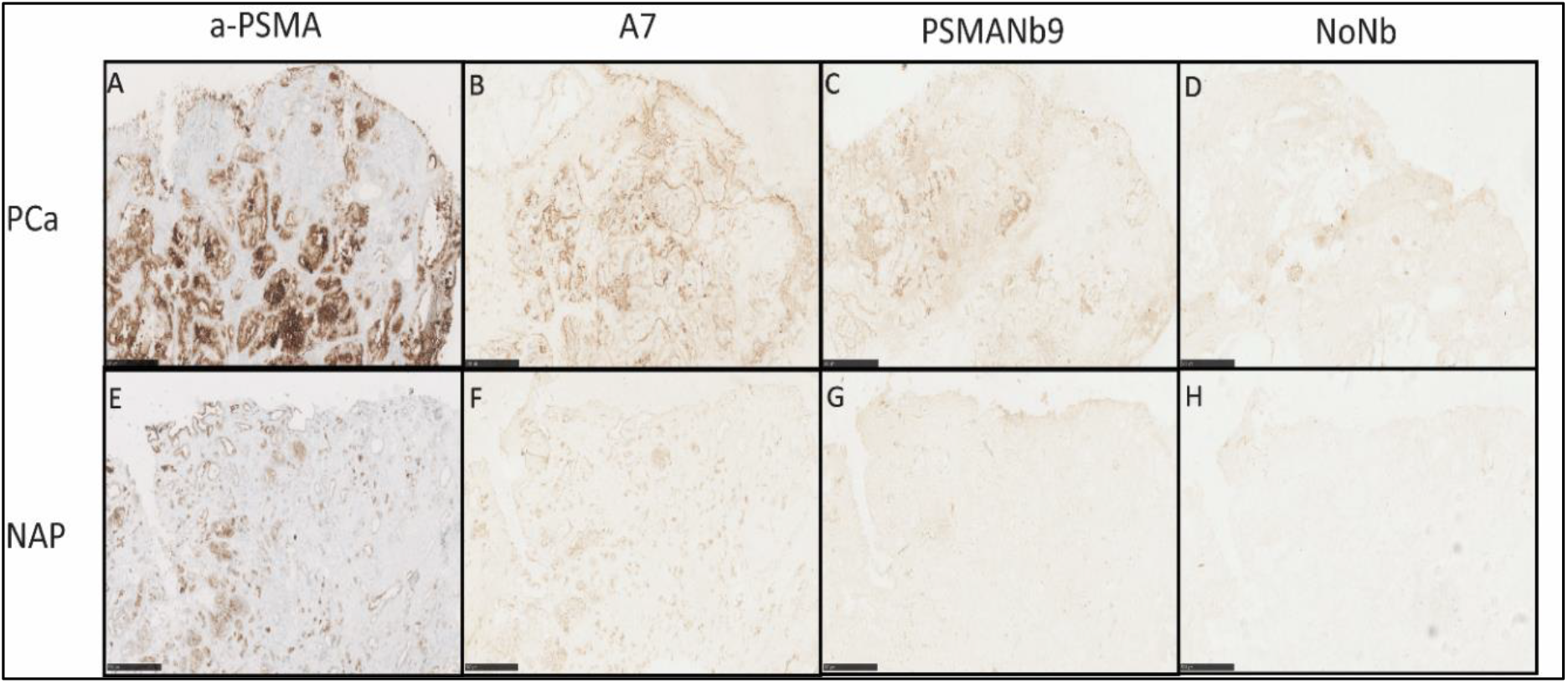
Fresh-frozen PCa (**A-D**) and normal adjacent prostate (NAP) (**E-H**) tissues from radical prostatectomy samples were cryo-sectioned and stained with the anti-PSMA antibody (a-PSMA) (**A, E**), A7-phage (**B, F**), PSMANb9-phage (**C, G**) and the phage without nanobody (NoNb) (**D, H**). HRP-conjugated secondary antibodies were used for DAB staining. Representative images are shown. Black bar represents 0.5 mm.

Staining of human normal frozen tissue microarrays (TMA) (Suppl. Figure 4) showed weak staining with the phage without a nanobody (NoNb), particularly in liver and small intestine tissues, indicating non-specific interactions of phage proteins with tissue proteins. In concordance, the positive control anti-CD9 antibody and anti-CD9 nanobody-phage (H6) (Suppl. Figure 5) bound strongly across most tissues (www.proteinatlas.org/ENSG00000010278-CD9/tissue). The anti-PSMA IgG antibody strongly bound to small intestine and to a lesser extent to salivary gland, kidney, and spleen, which was in accordance with the levels of PSMA expression known for these tissues (www.proteinatlas.org/ENSG00000086205-FOLH1/tissue). Similarly, PSMANb9 and A7 nanobody phages demonstrated binding to small intestine, salivary glands and kidneys with an overall weaker binding across other normal tissues, which could be attributed to the non-specific interactions of phage coat proteins.

#### 2.7 A7 and PSMANb9 nanobody-phages bind PSMA protein on the surface of PSMA positive cell lines

To show that nanobody-phages A7 and PSMANb9 interact with the PSMA protein on the surface of PSMA positive cell lines, PSMA expression was knocked down in LNCaP cells using shRNA technology as confirmed by Western blot analysis where a significant knockdown of PSMA was observed in LNCaP-shPSMA cells compared to LNCaP-WT and LNCaP-shNC (negative control shRNA) cells (Figure 4A). Next, we evaluated the binding of ATTO488-labeled nanobody-phages A7 and PSMANb9 to LNCaP-WT and LNCaP-shPSMA cells using flow cytometry. A sharp reduction in anti-PSMA antibody and nanobody signals were observed when bound to LNCaP-shPSMA cells compared to LNCaP-WT and LNCaP-shNC cells (Figure 4B).

**Figure 4:**
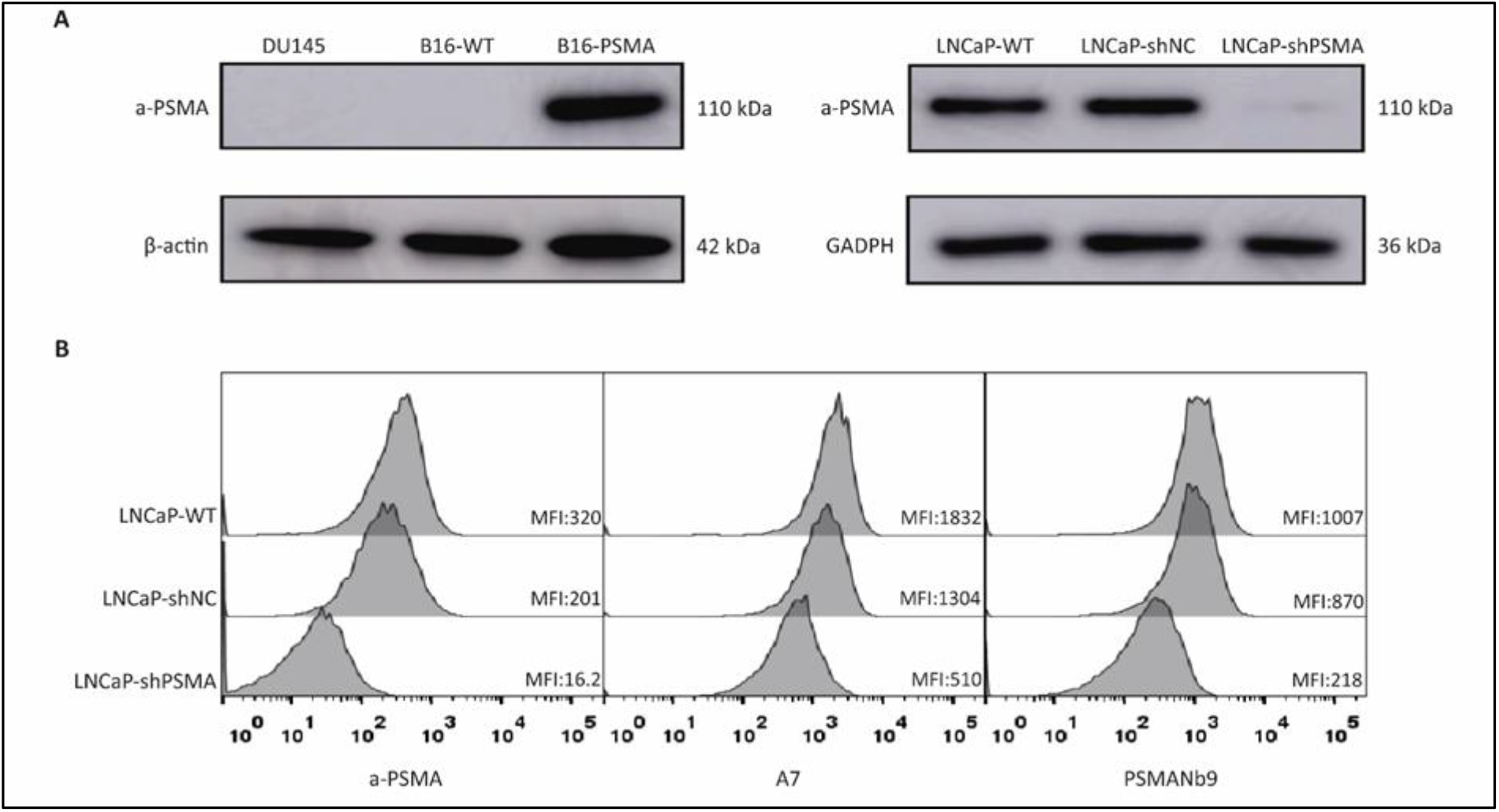
**(A)** PSMA expression in PSMA positive and negative cell lines in Western blots using anti-PSMA (a-PSMA) antibody and anti β-actin and GAPDH as loading controls. **(B)** Flow cytometry of binding of a-PSMA antibody, ATTO488-labeled nanobody-phages A7 and PSMANb9 to LNCaP cells. MFI: Median Fluorescence Intensity.

#### 2.8 Computational modeling of nanobody-PSMA interactions predicts two distinct epitopes for A7 and PSMANb9

The 3D structures of candidate nanobodies were modeled using two approaches: the homology-based modeling tool SWISS-MODEL [25] and the RoseTTAFold [26] option of the Robetta server, a deep learning-based modeling platform that also incorporates *de novo* structure prediction for domains lacking sequence homology. The generated models were evaluated comprehensively using ERRAT scores, MolProbity scores, PROCHECK residue statistics, and ProSA-web Z-scores to assess their structural reliability (Suppl. Table 2).

Overall, both Robetta and SWISS-MODEL produced nanobody models of good structural quality. Robetta models generally demonstrated higher ERRAT scores and more favorable ProSA-web Z-scores, reflecting better non-bonded atomic interactions and overall fold quality. PROCHECK analysis showed that the majority of models were well-refined, with more than 90% of residues located within favored and allowed regions of the Ramachandran plot, indicating proper backbone geometry (Suppl. Table 2). ProSA-web results supported the high quality of all models, as the Z-scores (≤ –6.03) were consistent with those reported for experimentally determined protein structures.

Despite the strong performance of both tools, Robetta-generated models were selected for further analysis due to their slightly superior scoring metrics. Moreover, Robetta’s ability to intergrate *de novo* prediction methods for regions without suitable templates provides an added advantage in modeling challenging nanobody domains.

The nanobody models were docked to the PSMA ectodomain structure (PDB ID: 1Z8L) using HADDOCK 2.4 to predict the interaction interface between each nanobody and PSMA ectodomain [27]. The docking generated multiple clusters of models ranked based on the HADDOCK score, which integrates van der Waals, electrostatic, desolvation energies, and restraint violations to evaluate binding affinity and interface quality. The best-ranked complexes exhibited favorable HADDOCK scores ranging from –120 to –83, indicating strong and reliable binding interactions. After obtaining the docked complexes from HADDOCK 2.4, the models were further analyzed using the PDBsum generator to identify specific interactions at the nanobody-PSMA interface. Based on the detailed analysis of hydrogen bonds (Suppl. Table 3), nanobodies A7, A3, PSMANb6, and PSMANb5 were found to recognize one distinct epitope on the PSMA ectodomain. In contrast, nanobodies PSMANb9 and PSMANb11 targeted a separate and non-overlapping region of PSMA. The PSMA-617 tracer binds a unique epitope close to the active glutamate carboxypeptidase site of PSMA, consistent with the crystallographic structure of PSMA-PSMA-617 (PDB ID: 505U) complex (Figure 5).

**Figure 5:**
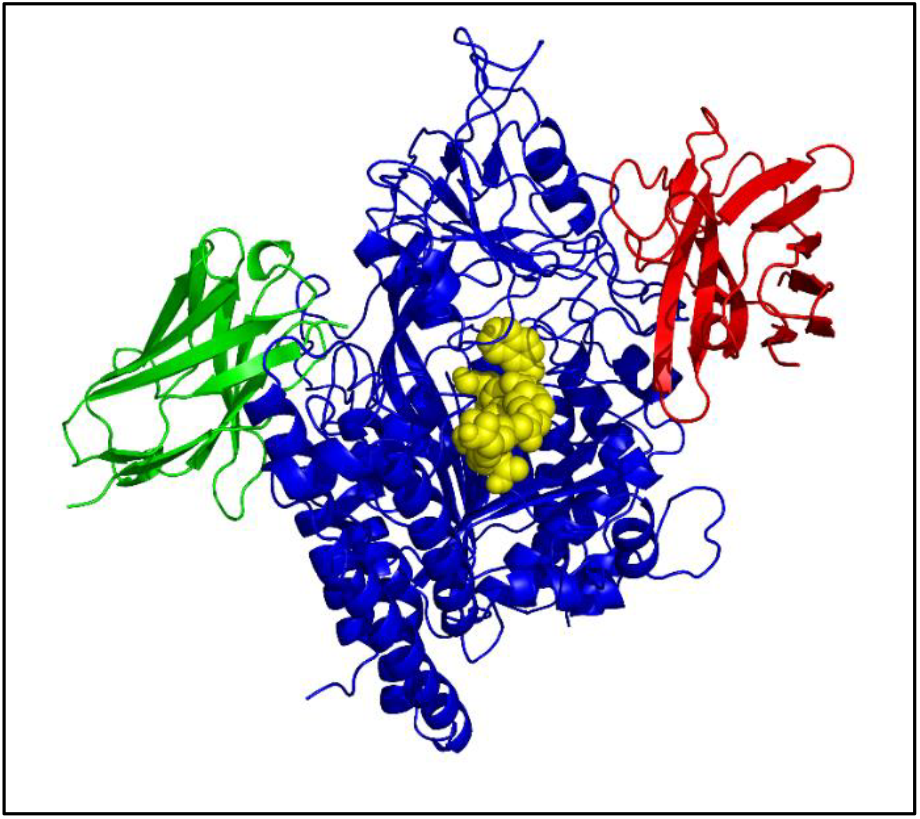
HADDOCK docking models of nanobody-PSMA complexes illustrating the interaction interface between PSMA ectodomain, two representative nanobodies and a tracer. The model reveals the different epitopes on PSMA (dark blue) for PSMANb9 (red), A7 (green) and the PSMA-617 tracer (yellow).

### 2.9 Molecular docking simulations indicate the high stability of nanobody-PSMA complexes

To assess the structural stability of PSMA-A7 and PSMA-PSMANb9 complexes, we performed 3 times 200 ns production MD (Molecular Dynamics) using GROMACS 2022 and OPLS-AA force field and examined Root Mean Square Deviation (RMSD), Radius of Gyration (Rg) and Root Mean Square Fluctuation (RMSF) parameters for each complex (Suppl. Figure 6). Both systems reached equilibrium after approximately 50–80 ns, but the PSMANb9-PSMA complex exhibited lower RMSD values (around 0.2–0.4 nm) compared to A7-PSMA (0.6–0.8 nm), indicating a more stable overall conformation (Suppl. Figure 6A). The radius of gyration (Rg), which reflects the overall molecular size and compactness, remained nearly constant for both complexes throughout the simulation. A slightly lower Rg observed for PSMANb9-PSMA suggests a more compact and stable conformation compared to A7-PSMA (Suppl. Figure 6B). Additionally, analyzing the fluctuations of backbone atoms in each residue showed slight fluctuations for the majority of residues, represented by RMSF values plotted against residue number (Suppl. Figure 6C, 6D). Residue-based RMSF analysis further revealed that PSMANb9-PSMA showed reduced local flexibility, particularly at the interface regions, supporting the observation of its enhanced structural stability. Together, these results suggest that the PSMANb9 forms a more stable and compact interaction with PSMA than A7.

### 2.10 PSMA docking predictions for binding of other selected nanobodies from the top 52 candidates

We tested whether there might be more than 2 predicted epitopes on PSMA when we would expand the modeling using the other non-tested 52 PSMA nanobodies identified earlier (Figure 1). A set of an additional 21 nanobodies with very different CDR3 sequences were selected for Robetta folding and HADDOCK docking. All of these nanobodies were predicted to bind to epitopes bound by A7 or PSMANb9 and no additional epitopes were identified (Suppl. Figure 7).

### 2.11 A7 and PSMANb9 nanobodies do not compete for PSMA binding

To validate the prediction of distinct epitopes on PSMA, we assessed the competitive binding of A7 and PSMANb9 nanobodies to LNCaP and B16-PSMA cell lines using a combination of nanobody proteins and nanobody-phages in a flow cytometry assay. The optimal concentration of the detector ATTO488-conjugated A7 and PSMANb9 nanobody-phages was determined using a concentration series in flow cytometry to LNCaP binding (Suppl. Figure 8). Concentrations of 5E+9 colony forming units (CFU) and 1E+10 CFU for ATTO488-labeled A7 and PSMANb9 nanobody-phages were selected, respectively. As unlabeled competitors, PSMANb9 and A7 His-tagged proteins were produced (Suppl. Figure 9). Based on the titration assay with increasing concentration of either A7 or PSMANb9 proteins we selected the 1 mg/mL concentration to saturate cell surface PSMA proteins (Suppl. Figure 10).

After blocking cell surface PSMA with A7 purified monovalent proteins in either LNCaP and B16-PSMA cells, the binding intensity of ATTO488-labeled A7 nanobody-phages dramatically decreased (Figure 6). This was similar for the PSMANb9 combination indicating competition of the nanobody protein and corresponding nanobody-phage for the same epitope. In contrast, when cells were blocked either with A7 protein before ATTO488-labeled PSMANb9-phage or PSMANb9 protein before ATTO488 labeled A7-phage, no significant change was observed in ATTO488 fluorescent signal intensity. This observation revealed that A7 and PSMANb9 bind to two distinct epitopes on the extracellular domain of PSMA, confirming the *in silico* prediction of two different epitopes of the PSMA protein (Figure 5).

**Figure 6:**
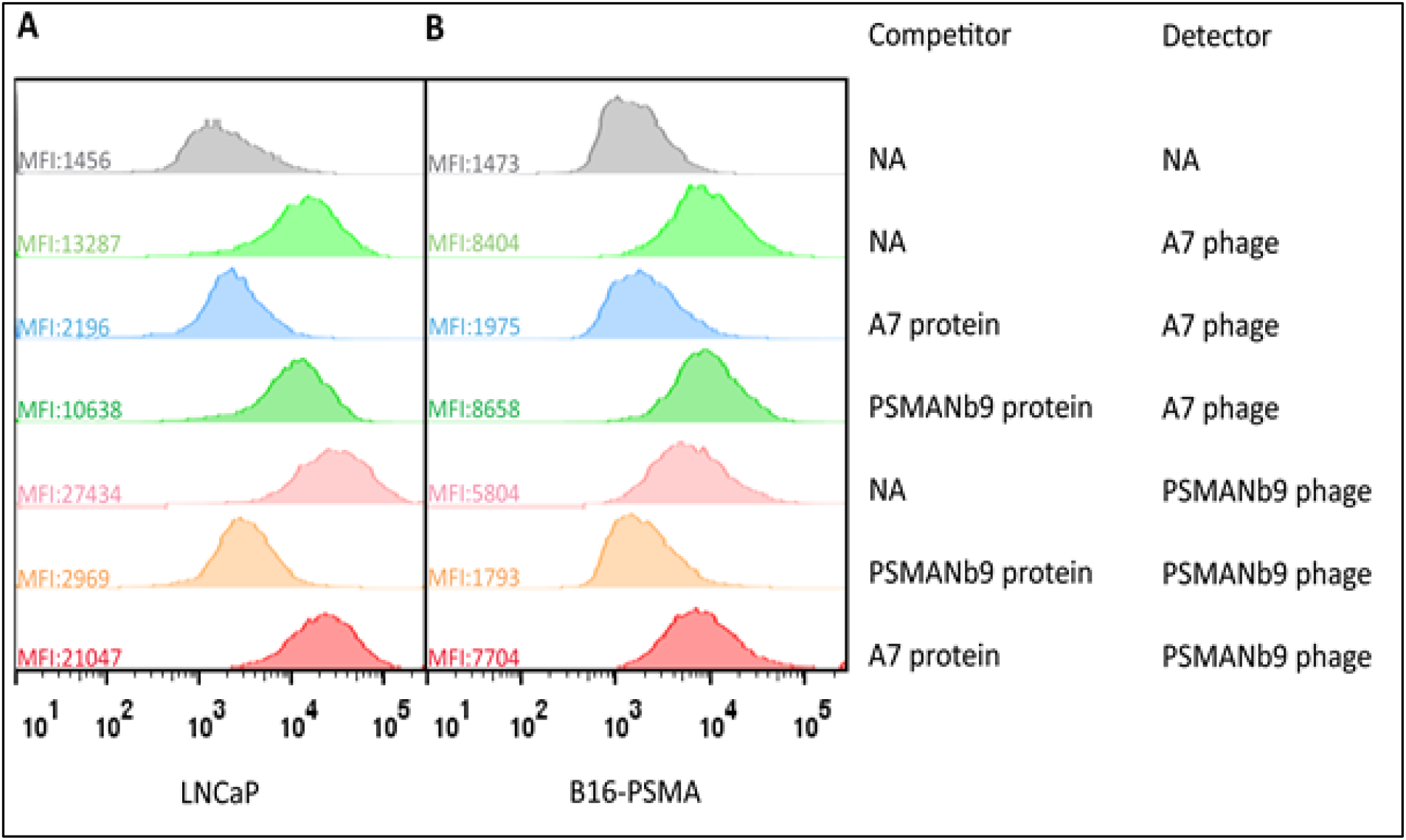
Epitope competition assay using A7 or PSMANb9 proteins with or without ATTO488-labeled A7-phage or PSMANb9-phage on LNCaP **(A)** and B16-PSMA **(B)** cells. Flow cytometry was performed by first a 1 hr incubation with purified A7 or PSMANb9 protein, and without any washing, followed by a 1 hr incubation with the ATTO488-labeled nanobody-phages. MFI: Median Fluorescence Intensity; NA, Not Available.

## 3. Discussion

### 3.1 Panning against collections of cell lines allows for identification of PSMA nanobodies

In the present study, we questioned whether non-targeted immunization using tumor cells and non-targeted panning using a large panel of cell lines with diverse protein expression profiles, could identify nanobodies against PSMA. Comparing nanobody binding to 4 PSMApos cell lines with 8 or 13 PSMAneg cell lines, allowed selection of 498 nanobody clusters preferentially binding PSMA positive cell lines. From these, 10% (n=52) were also preferentially binding the B16-PSMA cell line over B16-WT, the targeted approach for selecting PSMA nanobodies (Figure 1C). The majority of the nanobodies from the panning of the cell line panel, does not seem to bind PSMA, which could be explained by potential binding to cell surface proteins co-expressed with PSMA such as STEAP1, PSCA, SLC44A4 and SLC39A6 [28]. In addition, any single round panning is expected to have a high number of non-specific binders that will not be fully eliminated when comparing two groups with limited numbers of cell lines.

### 3.2 Multiple rounds negative-positive panning as compared to single round panning

Whether the classical 3 round negative-positive panning (3R-NegPos) outperforms the 1 round single cell pannings (1R-SC), was tested by comparing 3R-NegPos and 1R-SC using the B16-WT and B16-PSMA cell lines. Only 6 nanobody clusters overlapped between the 462 and 341 clusters (Figure 1A).

Multiple factors underlie this limited overlap. First, nanobody clusters enriched after 1R-SC were more prevalent in the original library, with an average count of 1145, compared to just 164 for the 462 3R-NegPos clusters. Second, repeated panning rounds likely amplify also nanobodies binding to minor surface protein differences between B16-PSMA and B16-WT, other than PSMA. Third, every additional panning round introduces a bias in clone amplification and loss of diversity [29]. Fourth, the sequencing depth, PCR amplification biases and nanobody selection criteria are expected to affect differences identified [30]. The strong enrichment for 6 overlapping nanobody clusters during the 3R-NegPos panning (up to 29.9%) resulted in their identification during conventional colony-picking [12]. This would be less likely using 1R-SC with only up to 0.41% enrichment of the 6 clusters.

### 3.3 Panning against collections of cell lines provides a powerful means of selection with restrictions

The use of a panel of cell lines with diverse protein expression such as a strong distinction in PSMA expression, allows for the selection of PSMA nanobodies upon untargeted single round panning. This approach comes with some limitations. First, besides endogenous cell line protein expression infections can result in differential nanobody binding. The Xenotropic Murine leukemia virus-Related Virus (XMRV) [31] is present in cell lines created through mouse engrafting. Except for LNCaP and MSKPCa12D, these unfortunately include all PSMA-positive PCa cell lines (PC346C, MDAPCa2b, 22Rv1, VCaP) and one PSMA-negative line (PC339C). Based on the binding of the XMRV-recognizing nanobodies, all other cell lines in the panel are XMRV-negative except for the mouse B16-WT and B16-PSMA. Their binding potentially reflects infection by the closely related MelARV or Bxv-1 viruses [32]. During the 3R-NegPos B16-WT and B16-PSMA panning, the XMRV binding nanobodies were selected out, which was not the case in the 1R-SC cell line pannings for which we corrected the *in silico* selections as described in Materials and Methods. The inventory and impact of viral infections of cell cultures are understudied and will distort the selection of cancer nanobodies if not taken into account [33]. Second, to determine PSMA positivity, RNA expression levels were used to separate cell lines into two groups. However, in VCaP cells, high FOLH1 mRNA and PSMA protein (by Western blotting) [34] did not correlate with strong antibody or nanobody binding, suggesting limited PSMA surface exposure. Inspection of RNA sequencing data, this discrepancy is not explained by PSMA splice variants or mutations in VCaP cells. It does highlights that RNA levels do not always reliably represent surface protein abundance [35]. Third, to confidently identify nanobodies, the protein of interest must show distinct expression across a large group of high and low expressing cell lines. Within our selection using 4 PSMApos cell lines and 8 or 13 PSMAneg cell lines, approximately 10% of the 498 nanobody clusters are PSMA binders based on overlay with the B16-PSMA targeted panning indicating limitations in PSMA-specific enrichment.

### 3.4 Validation and prediction of distinct PSMA binding epitopes and utilization of docking software

Of the 12 selected nanobodies tested on cell-ELISA, 6 did not significantly reproduce the panning data and were discarded for further validation. Both the ELISA and IHC tests were performed using the easy-to-use nanobody-phage format, which resulted in higher background as compared to purified antibodies or nanobodies and could explain some of the variable reproducibility. Computational modeling and docking simulations using the validated nanobodies revealed two distinct PSMA epitopes, which were verified through epitope competition assays between nanobodies A7 and PSMANb9. While homology modeling using Robetta and SWISS-MODEL provided comparable structural predictions, accurate modeling of the diverse CDR3 loop remains challenging due to lack of sufficient sequence homology [36]. Although Robetta uses *ab initio* modeling for loop regions, no significant difference was observed between quality indicators (Supp Table 2). Leveraging deep learning tools like H3-OPT [37] and DeepH3 [37] offers promising alternatives.

HADDOCK-guided docking showed that the majority of nanobody interactions occur via CDR3, with no evidence of a third PSMA epitope among 27 candidate nanobodies. However, the exact interacting residues should be further verified through standard methods [38, 39].

### 3.5 Future perspectives

Based on the observation that A7 and PSMANb9 nanobodies bind different epitopes, allows for the development of novel biparatopic tracers to increase affinity and internalization due to receptor surface clustering [40]. Such molecules could further expand the exploitation of PSMA in diagnostics, PET/SPECT and intraoperative imaging and therapeutics [41].

Cellular immunization and non-targeted panning against a panel of cell lines could be the basis of identification of cell surface targets. The classification of cell lines into groups with and without PSMA expression allowed for the selection of nanobodies potentially binding this PCa target. At the moment, such a selection alone is not powerful enough to have almost certainty of PSMA binding and a targeted approach using B16-WT and B16-PSMA panning was needed to enrich for PSMA binders. Having access to the increasing number of nanobody sequences and their target proteins, fuels research on VHH structural modelling and docking [21, 22]. The next steps towards AI-driven target prediction and *de novo* nanobody design are being taken and will strongly benefit from large panning databases [23, 24].

## 4. Materials and Methods

Details of the various protocols can be found in the Supplementary information.

### 4.1 Llamas immunization and VHH-library construction

Three Llamas were immunized with four PCa cell lines (LNCaP, PC346C, VCaP, and MDAPCa2b) and a combination of fresh frozen PCa and bladder cancer (BlCa) cell lines, from which three nanobody-phage libraries were constructed. Characteristics of the three different libraries are presented in Table 1.

### 4.2 Screening and selection of nanobodies using phage display technology

To produce phage display libraries, each nanobody library was infected with M13 K07Δplll hyperphages (Progen Biotechnik, Germany). To screen PSMA-binding nanobodies, the full L1P4 library or the mix of three libraries were used in a 0.13:0.52:0.35 ratio and screened using three biopanning strategies: three rounds of negative-positive panning (3R-NegPos) against B16-WT and B16-PSMA, one round of single cell panning (1R-SC) against B16-WT and B16-PSMA and one round of biopanning against PSMA positive cells and PSMA negative cells cancer cell lines (1R-Cell Line Panning). The biopanning protocol was adapted from Chatalic et al. [12]. For 3R-NegPos, each round consisted of a negative selection against B16-WT cells using 100x the L1P4 library followed by a positive selection against B16-PSMA cells. In the 1R-SC strategy, one individual positive selection was performed against B16-WT and B16-PSMA, respectively. For the 1R-Cell line panning, one single positive selection was performed on each cell line, separately.

To select B16-PSMA specific nanobodies, the conventional colony picking method was described previously [12]. For the NGS analysis of the selected phages at the end of each panning round, the nanobody cDNA from each nanobody-phage sample corresponding to an individual cell line from a single selection round was amplified and sequenced using the Illumina MiSeq 2x300 bp protocol to a 1 mln paired read-depth per sample (Illumina MiSeq Reagent Kits v3). For each sample and panning round, a specific barcode (index) was present in the PCR amplification primers to allow multiplexing of samples in each MiSeq run. NGS data was processed using an in-house developed pipeline described in the Supplementary information.

### 4.3 Selection of PSMA-binding nanobodies from nanobody database

To select PSMA-binding nanobodies, we defined certain criteria within each biopanning strategy. For 3R-NegPos, we first determined the ratio between selection round (SR) 2/1 and 3/1 with respect to the frequency of each unique nanobody sequence and selected candidate nanobodies for which both the SR2/SR1 and SR3/1 was greater than 5, while the value in SR3 was set to greater than 1000 (is 0.001% of scaled 100 million). For 1R-SC, the ratio between B16-PSMA and B16-WT should be greater than 5, and the value of the average B16-PSMA greater than 1000. For the 1R-Cell Line Panning, the average of nanobody binding to PSMApos and PSMAneg cells was determined where nanobodies with a PSMApos/PSMAneg ratio greater than 5, XMRVpos/XMRVneg ratio less than 3 and average PSMApos greater than 1000 were selected as top candidates.

The DNA sequence of each selected nanobody was retrieved from sequencing data to produce nanobody-phages which were further characterized.

### 4.4 Hierarchical clustering and circular dendrogram of nanobody CDR3 sequences

Nanobody CDR3 sequences were clustered based on pairwise Levenshtein distances using average-linkage hierarchical clustering. The resulting dendrogram was displayed in a circular configuration using R (igraph, ggraph, stringdist). Branches were colored by dominant library—L1P4 (coral red), LUPCa1 (green), and LUPCa2 (blue)—with progressively darker shades indicating deeper hierarchical levels. Selected nanobodies were indicated by black outer nodes, whereas unselected ones were represented by light gray.

### 4.5 Whole cell-ELISA

PSMA binding of the selected nanobodies was assessed by cell-ELISA. The experiment was performed using two PSMA-positive cell lines, LNCaP and B16-PSMA, along with two PSMA-negative cell lines, DU145 and B16-WT. Nanobody binding was detected using mouse anti-M13 antibody as the primary antibody followed by goat anti-mouse HRP secondary antibody and OPD substrate. Antibody information is provided in Suppl. Table 5.

### 4.6 Cell immunohistochemistry (IHC)

To investigate the specific binding of nanobodies, we created a cell array where 20 cell lines were coated on a glass cavity slide (Marienfeld, Germany) and stained with nanobodies. Briefly, cells were blocked with 1 ml 1% BSA-PBS and then stained with 5E+10 PFU/slide of nanobody-phages. Nanobody binding was detected using mouse anti-M13 antibody followed by goat anti-mouse HRP secondary antibody and DAB substrate.

### 4.7 Immunohistochemistry staining of frozen tissue microarrays and frozen prostate tissue sections

Tissue microarray (TMA) slides representing 20 human normal tissues in duplicates were purchased from Biochain, USA. Frozen human prostate normal and cancer tissues were obtained from the Erasmus MC Tissue Biobank (Rotterdam, Netherlands), sectioned with 5-10 um thickness and attached to Starfrost Adhesive slides (Deltalab, Spain) and frozen in −80°C. Immunohistochemistry (IHC) staining of tissues was performed according to the protocol described above. Slides were scanned by PARTS (Erasmus MC Pathology and Research and Trial Service, Rotterdam, The Netherlands) using Nanozoomer 2.0 HT digital slide scanner (Hamamatsu Photonics K.K, Germany) and images were visualized and analyzed using NDP viewer software

### 4.8 Lentiviral Transduction of shRNA for PSMA Knockdown

HEK293T cells were transfected using calcium phosphate transfection kit with a combination of 5 different PSMA shRNAs to produce lentiviruses (Mission shRNA libraries Sigma-Aldrich, Center for Biomics, Erasmus MC). A GFP-Luciferase fusion construct (M21) and non-targeting human gene shRNA (shNC) served as positive and negative control, respectively. The virus containing supernatant were collected 48 hrs post-transfection and used to infect LNCaP cells.

### 4.9 Western blot analysis of PSMA expression

Cells were lysed using RIPA buffer and total protein was extracted. Five µg of each protein sample including wild type (WT), knockdown (shPSMA) and shRNA negative control (shNC) LNCaP cells along with DU145, B16-WT and B16-PSMA were resolved on a 10% SDS-PAGE gel and transferred to a PVDF membrane. PSMA was detected using rabbit monoclonal anti-PSMA antibody while anti-β-actin antibody and anti-human GAPDH antibody were used as controls. Protein bands were detected using goat-anti-rabbit HRP or goat-anti-mouse HRP followed by BM chemiluminescence blotting substrate.

### 4.10 Binding of nanobodies to LNCaP cells in a flow cytometry assay

Binding of nanobody-phages to wild type (WT) and PSMA knockdown (shPSMA) LNCaP cells were investigated in a flow cytometry assay. For detection, A7 and PSMANb9 nanobody-phages were directly labeled with 0.25 mM ATTO488 NHS-ester dye (ATTO-TEC, Germany). Mouse monoclonal anti-PSMA antibody and goat-anti-mouse IgG cross-adsorbed secondary antibody Alexa Fluor 647 were used to detect PSMA expression in LNCaP-WT and shPSMA LNCaP cells.

### 4.11 Nanobody protein production

A7 and PSMANb9 nanobodies were cloned into the pHEN vector (Twist Bioscience, US) and proteins expressed including a C-terminal Myc-tag and 6xHis tag using *E. coli* HB2151 cells as host. Nanobody proteins were purified on a Ni-NTA agarose column using standard 25-500 mM imidazole gradient.

### 4.12 *In silico* modeling and docking of nanobody-PSMA complex

The 3D structure of human PSMA ectodomain (PDB ID: 1Z8L) was processed using BIOVIA Discovery Studio Visualizer to remove water molecules and unbound atoms. The 3D structures of candidate nanobodies-A3, A7, PSMANb5, PSMANb6, PSMANb9 and PSMANb11, were predicted using Robetta (http://robetta.bakerlab.org) and SWISS-MODEL server (https://swissmodel.expasy.org) both of which leverage homology modeling method [26]. To generate more accurate models, particularly within CDR3 regions, we used RoseTTAFold, a software tool that employs deep learning methods for accurate prediction of protein structures [26]. The predicted models were downloaded and evaluated for their overall quality (Suppl. Information).

Nanobody models and PSMA ectodomain were prepared for docking simulation using Dock Prep tool in UCSF Chimera followed by docking simulation of nanobody-PSMA monomer using the “easy interface” of HADDOCK 2.4 (High Ambiguity Driven-protein DOCKing) web server (https://wenmr.science.uu.nl/haddock2.4). The generated models were clustered and ranked based on their HADDOCK score. To compare the PSMA epitopes bound by PSMA-617 and our selected nanobodies, we also docked PSMA617 peptide to PSMA monomer using PyRx 0.8 [42] and Autodock 4.2 (The Scripps Research Institute, La Jolla, CA, USA) [43]. Molecular interactions were visualized and analyzed using Python Molecular Graphics (PyMOL, Schrödinger, LLC, USA) and LigPlot+ software [44]. The stability of nanobody-PSMA complex models was evaluated using GROMACS 2022 package (GROningen MAchine for Chemical Simulations, the Netherlands) and OPLS-AA force field (Optimized Potential for Liquid Simulations-All Atom, Yale University, USA) [45].

### 4.13 Epitope binding competition assay

A titration assay was conducted to determine the nanobody protein concentration sufficient to fully block the PSMA binding sites on the cell surface. LNCaP and B16-PSMA cells were incubated with varying concentrations of A7 or PSMANb9 nanobody proteins and binding was detected using a mouse monoclonal anti-Myc tag antibody followed by a goat anti-mouse Alexa Fluor 647 secondary antibody. Similar titration was performed using varying concentrations of ATTO488-labeled A7 or PSMANb9 nanobody-phages. For competition, LNCaP cells or B16-PSMA were first incubated with A7 or PSMANb9 proteins and without washing, followed by adding ATTO488-labeled PSMANb9 or A7 nanobody-phages.

### 4.14 Statistics

For hierarchical clustering and visualization and statistical analyses we used GraphPad Prism 9 (San Diego, USA). The hypergeometric Test was used for determining p-values of overlap in the Venn diagrams. The nonparametric Mann-Whitney test was applied for the cell-ELISA.

## Author contributions

TY performed the validation assays. JVZ cultured cells and performed the nanobody panning. ML performed pannings against 40 cell lines. SE and MVvdB cultured cells and processed clinical samples for immunization. RJ, AK and HvdW performed and supervised the data processing. PC created the L1P4 library. EB performed Miseq sequencing. CHB and NL supervised and provided input in the clinical aspects. AK and JVZ characterized the H6 nanobody. JL provided essential input in the panning concept. WMW supervised and was responsible for the A3, A5 and A7 nanobody identification. SK with assistance of KA performed the folding and docking. RT and GJ designed and supervised the research. TY, SK, GJ generated the figures and tables. TY, RT, SK and GJ wrote the manuscript. All authors edited the manuscript and approved the final version.

## Acknowledgements

The work was made possible through the “IMMPROVE” consortium sponsored by an Alpe d’Huzes grant of the Dutch Cancer Society (grant #EMCR2015-8022), through the projects CCBC and ‘Bladder cancer nanobodies’ from the Erasmus MC Daniel den Hoed Foundation and through the Chinese Scholarship Council program (grant number 202207650046).

We would like to thank Martijn van Duijn, Yann Seimbille and Erik Verburg for helpful discussions and Vincent Roodzant and Caitlin Jenster for assistance in tissue culture and pannings. We would like to thank Peter Riegman (department of Pathology) as head of the Erasmus MC Tissue Biobank. Cell lines were collected within the Erasmus MC and we thank all researchers making them available.

## Supporting information

Additional supporting information can be found in the Supplementary Information document.

## References

1. Smith, G.P., Filamentous fusion phage: novel expression vectors that display cloned antigens on the virion surface. Science, 1985. 228(4705): p. 1315–7.

2. McCafferty, J., et al., Phage antibodies: filamentous phage displaying antibody variable domains. Nature, 1990. 348(6301): p. 552–4.

3. Hamers-Casterman, C., et al., Naturally occurring antibodies devoid of light chains. Nature, 1993. 363(6428): p. 446–8.

4. Alexander, E. and K.W. Leong, Discovery of nanobodies: a comprehensive review of their applications and potential over the past five years. Journal of Nanobiotechnology, 2024. 22(1): p. 661.

5. Deschaght, P., et al., Large Diversity of Functional Nanobodies from a Camelid Immune Library Revealed by an Alternative Analysis of Next-Generation Sequencing Data. Front Immunol, 2017. 8: p. 420.

6. Shooli, H., et al., Theranostics in Brain Tumors. PET Clin, 2021. 16(3): p. 397–418.

7. Afshar-Oromieh, A., et al., The diagnostic value of PET/CT imaging with the (68)Ga-labelled PSMA ligand HBED-CC in the diagnosis of recurrent prostate cancer. Eur J Nucl Med Mol Imaging, 2015. 42(2): p. 197–209.

8. Eder, M., et al., 68Ga-complex lipophilicity and the targeting property of a urea-based PSMA inhibitor for PET imaging. Bioconjug Chem, 2012. 23(4): p. 688–97.

9. Weineisen, M., et al., 68Ga- and 177Lu-Labeled PSMA I&T: Optimization of a PSMA-Targeted Theranostic Concept and First Proof-of-Concept Human Studies. J Nucl Med, 2015. 56(8): p. 1169–76.

10. Kratochwil, C., et al., 225Ac-PSMA-617 for PSMA-Targeted α-Radiation Therapy of Metastatic Castration-Resistant Prostate Cancer. J Nucl Med, 2016. 57(12): p. 1941–1944.

11. Kulkarni, H.R., et al., PSMA-Based Radioligand Therapy for Metastatic Castration-Resistant Prostate Cancer: The Bad Berka Experience Since 2013. J Nucl Med, 2016. 57(Suppl 3): p. 97S–104S.

12. Chatalic, K.L., et al., A Novel ^111^In-Labeled Anti-Prostate-Specific Membrane Antigen Nanobody for Targeted SPECT/CT Imaging of Prostate Cancer. J Nucl Med, 2015. 56(7): p. 1094–9.

13. Evazalipour, M., et al., Generation and characterization of nanobodies targeting PSMA for molecular imaging of prostate cancer. Contrast Media Mol Imaging, 2014. 9(3): p. 211–20.

14. Fan, X., et al., Ultrasonic Nanobubbles Carrying Anti-PSMA Nanobody: Construction and Application in Prostate Cancer-Targeted Imaging. PLoS One, 2015. 10(6): p. e0127419.

15. Hassani, M., et al., Construction of a chimeric antigen receptor bearing a nanobody against prostate a specific membrane antigen in prostate cancer. J Cell Biochem, 2019. 120(6): p. 10787–10795.

16. Nonnekens, J., et al., (213)Bi-Labeled Prostate-Specific Membrane Antigen-Targeting Agents Induce DNA Double-Strand Breaks in Prostate Cancer Xenografts. Cancer Biother Radiopharm, 2017. 32(2): p. 67–73.

17. Rosenfeld, L., et al., Nanobodies Targeting Prostate-Specific Membrane Antigen for the Imaging and Therapy of Prostate Cancer. J Med Chem, 2020. 63(14): p. 7601–7615.

18. Ruigrok, E.A.M., et al., Extensive preclinical evaluation of lutetium-177-labeled PSMA-specific tracers for prostate cancer radionuclide therapy. Eur J Nucl Med Mol Imaging, 2021. 48(5): p. 1339–1350.

19. Xing, Y., et al., A Single-Domain Antibody-Based Anti-PSMA Recombinant Immunotoxin Exhibits Specificity and Efficacy for Prostate Cancer Therapy. Int J Mol Sci, 2021. 22(11).

20. Su, J., et al., Nanobodies: a new potential for prostate cancer treatment. J Cancer Res Clin Oncol, 2023. 149(9): p. 6703–6710.

21. Mitchell, L.S. and L.J. Colwell, Comparative analysis of nanobody sequence and structure data. Proteins, 2018. 86(7): p. 697–706.

22. Vishwakarma, P., et al., V(H)H Structural Modelling Approaches: A Critical Review. Int J Mol Sci, 2022. 23(7).

23. El Salamouni, N.S., et al., Nanobody engineering: computational modelling and design for biomedical and therapeutic applications. FEBS Open Bio, 2025. 15(2): p. 236–253.

24. Zhu, H. and Y. Ding, Nanobodies: From Discovery to AI-Driven Design. Biology (Basel), 2025. 14(5).

25. Waterhouse, A., et al., SWISS-MODEL: homology modelling of protein structures and complexes. Nucleic acids research, 2018. 46(W1): p. W296–W303.

26. Baek, M., et al., Accurate prediction of protein structures and interactions using a three-track neural network. Science, 2021. 373(6557): p. 871–876.

27. Honorato, R.V., et al., The HADDOCK2. 4 web server for integrative modeling of biomolecular complexes. Nature protocols, 2024. 19(11): p. 3219–3241.

28. Sharifi, M.N., et al., Clinical cell-surface targets in metastatic and primary solid cancers. JCI insight, 2024. 9(18): p. e183674.

29. Derda, R., et al., Diversity of phage-displayed libraries of peptides during panning and amplification. Molecules, 2011. 16(2): p. 1776–1803.

30. Ac’t Hoen, P., et al., Phage display screening without repetitious selection rounds. Analytical biochemistry, 2012. 421(2): p. 622–631.

31. Hempel, H.A., et al., Infection of xenotransplanted human cell lines by murine retroviruses: a lesson brought back to light by XMRV. Frontiers in Oncology, 2013. 3: p. 156.

32. Li, M., et al., Sequence and insertion sites of murine melanoma-associated retrovirus. Journal of virology, 1999. 73(11): p. 9178–9186.

33. Uphoff, C.C., et al., Screening human cell lines for viral infections applying RNA-Seq data analysis. PLoS One, 2019. 14(1): p. e0210404.

34. Kuzmanov, A., et al., Regulation of prostate-specific membrane antigen (PSMA) expression in prostate cancer cells after treatment with dutasteride and lovastatin. Neoplasia, 2024. 57: p. 101045.

35. Nusinow, D.P., et al., Quantitative proteomics of the cancer cell line encyclopedia. Cell, 2020. 180(2): p. 387-402. e16.

36. Ruffolo, J.A., et al., Fast, accurate antibody structure prediction from deep learning on massive set of natural antibodies. Nature communications, 2023. 14(1): p. 2389.

37. Chen, H., et al., Accurate prediction of CDR-H3 loop structures of antibodies with deep learning. elife, 2024. 12: p. RP91512.

38. Islam, Z., et al., Structural insights into the unique recognition module between α-synuclein peptide and nanobody. Protein Science, 2024. 33(2): p. e4875.

39. Pleiner, T., et al., Nanobodies: site-specific labeling for super-resolution imaging, rapid epitope-mapping and native protein complex isolation. elife, 2015. 4: p. e11349.

40. Nessler, I., et al., Increased tumor penetration of single-domain antibody–drug conjugates improves in vivo efficacy in prostate cancer models. Cancer research, 2020. 80(6): p. 1268–1278.

41. Zeng, T., et al., The Application of Prostate Specific Membrane Antigen in the Diagnosis and Treatment of Prostate Cancer: Status and Challenge. OncoTargets and Therapy, 2024: p. 991–1015.

42. Dallakyan, S. and A.J. Olson, Small-molecule library screening by docking with PyRx, in Chemical biology: methods and protocols. 2014, Springer. p. 243–250.

43. Morris, G.M., et al., AutoDock4 and AutoDockTools4: Automated docking with selective receptor flexibility. Journal of computational chemistry, 2009. 30(16): p. 2785–2791.

44. Wallace, A.C., R.A. Laskowski, and J.M. Thornton, LIGPLOT: a program to generate schematic diagrams of protein-ligand interactions. Protein engineering, design and selection, 1995. 8(2): p. 127–134.

45. Jorgensen, W.L., D.S. Maxwell, and J. Tirado-Rives, Development and testing of the OPLS all-atom force field on conformational energetics and properties of organic liquids. Journal of the american chemical society, 1996. 118(45): p. 11225–11236.

