## Supplemental Information for "Cell-Based Immunization Combined with Single-Round Cell Panning Enables Discovery of PSMA-Targeting Nanobodies from Phage Display Libraries"

#### 1. Supplementary Figures

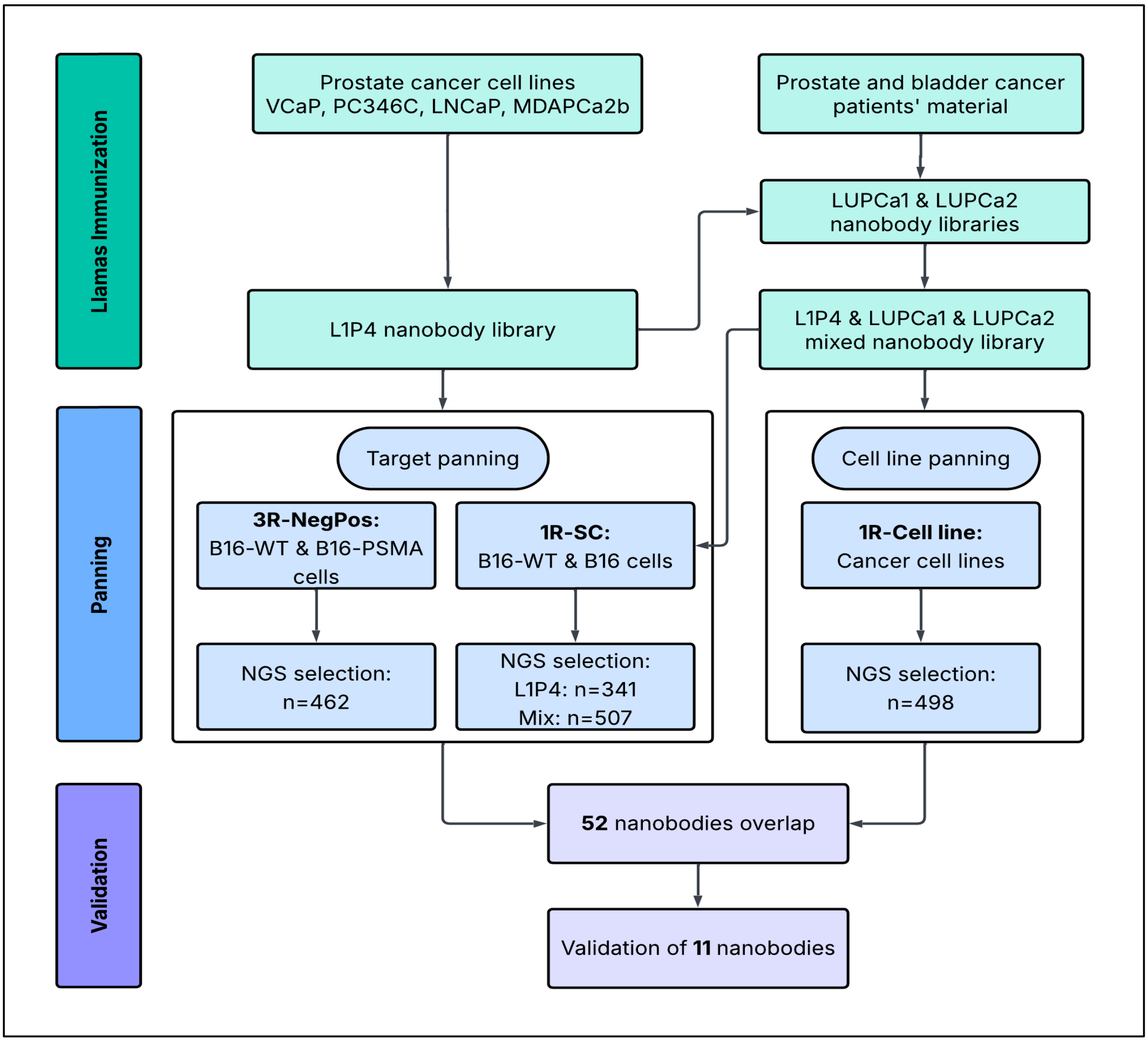


*Suppl. Figure 1: Overview of the panning and selection strategy to identify nanobodies against PSMA. Nanobody-phage libraries were created via Llama immunization using cell lines (L1P4) or cell fractions from patient tumor samples (LUPCa1 and LUPCa2). These libraries were used for panning against B16-WT and B16-PSMA in a 3-round negative-positive (3R-NegPos) or 1 round single cell (1R-SC) panning protocol. One round panning was also performed against a series of cell lines for which most do not express PSMA and a selection of the prostate cancer cell lines that do. Bound nanobody-phages were extracted and sequenced using Illumina Next Generation Sequencing (NGS) between 400,000-1,200,000 reads deep. Nanobodies with an identical CDR3 amino acid sequence were grouped into one cluster and the number of reads per panning counted. For each panning, nanobody clusters were selected that were more abundant in PSMA positive pannings and the overlap between pannings compared.*

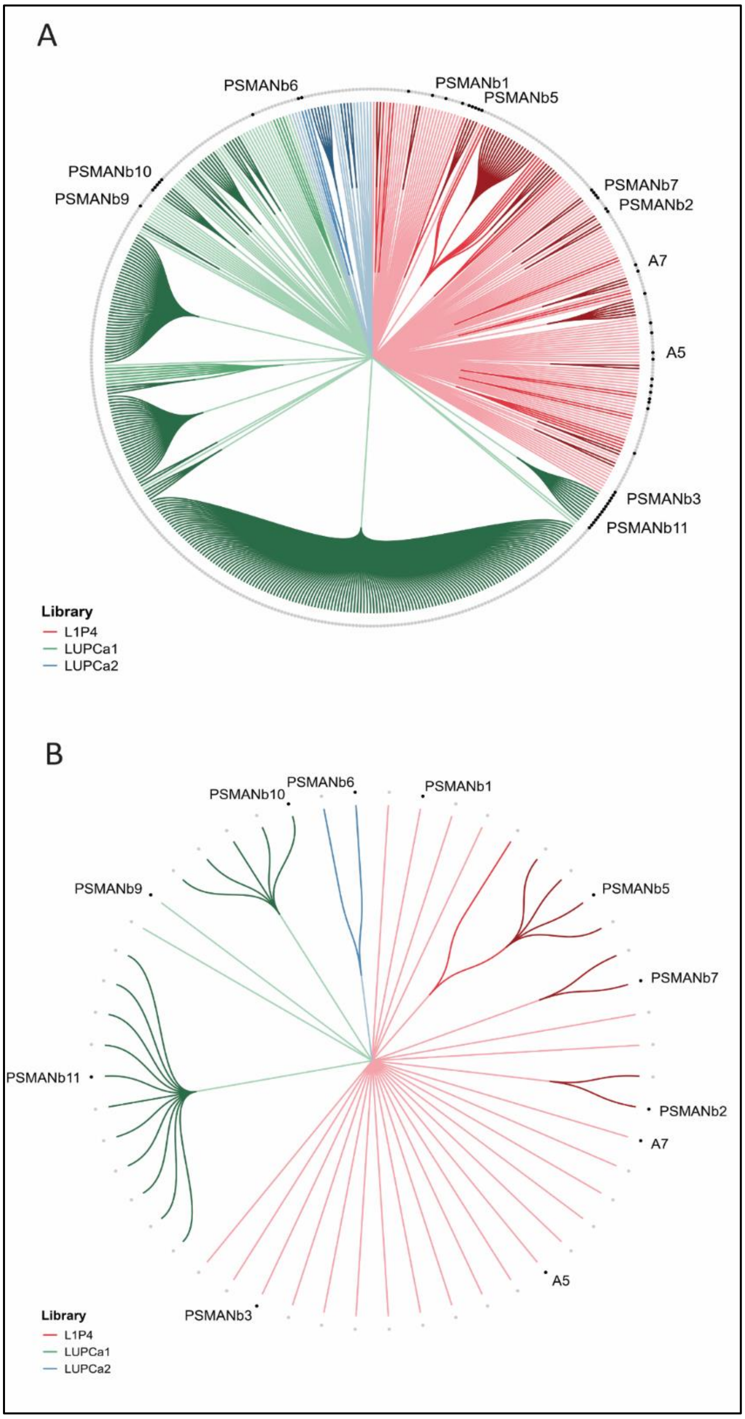


*Suppl. Figure 2: Circular dendrograms of nanobody CDR3 sequences.* ***(A)*** *Dendrogram of 498 nanobodies from 1R-SC PSMApos cell line panning.* ***(B)*** *Dendrogram of 52 candidate nanobodies selected from overlapping group (Figure 1C). Hierarchical clustering of nanobody CDR3 repertoires was visualized in a circular dendrogram. Branch colors represent the predominant library origin—L1P4 (coral red), LUPCa1 (green), and LUPCa2 (blue)—with light, medium, and dark shades denoting increasing hierarchical depth. Black outer nodes (****A****: 52 overlapping nanobodies;* ***B****: 11 PSMA nanobodies for validation) and gray outer nodes indicate selected and unselected nanobodies, respectively. Labeled clones for validation are annotated at a fixed radial distance.*

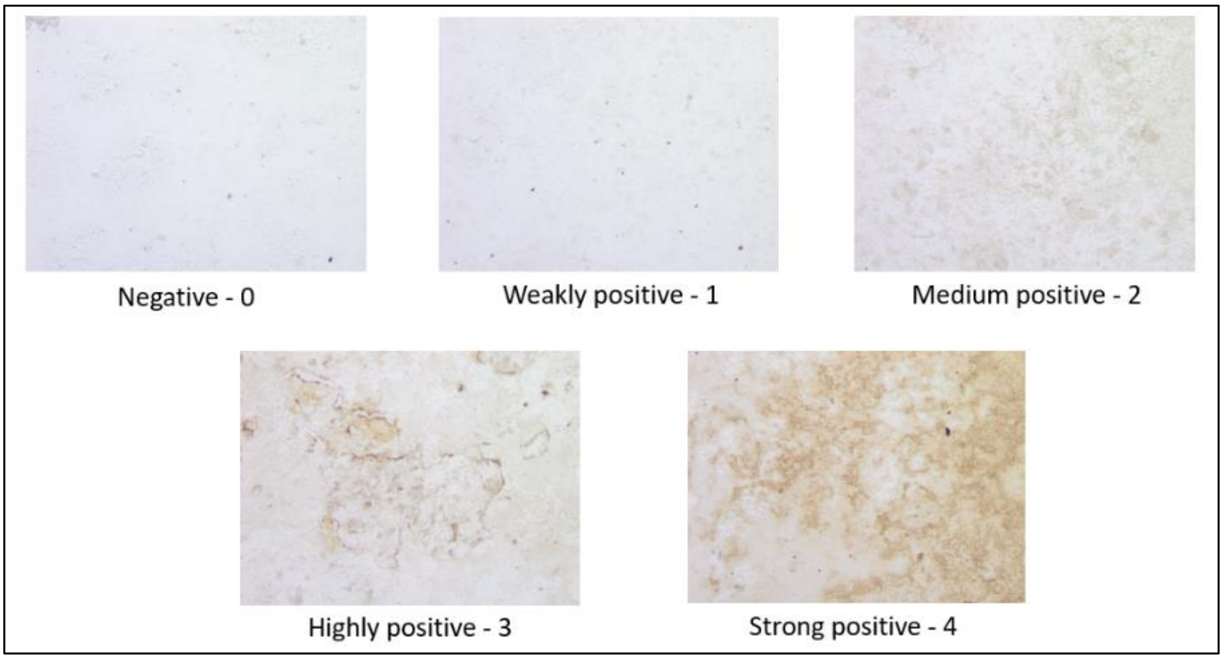


*Suppl. Figure 3: Representative images of various staining intensities indicating the scoring scheme with no staining scored as 0 to strongly stained cells scored as 4. Cells spotted on glass slides were incubated with nanobody-phages followed by an anti-M13-HRP antibody and DAB color staining. Two independent scientists blindly scored the staining intensities of nanobody-phages to 20 different cell lines using the 0-4 intensity system.*

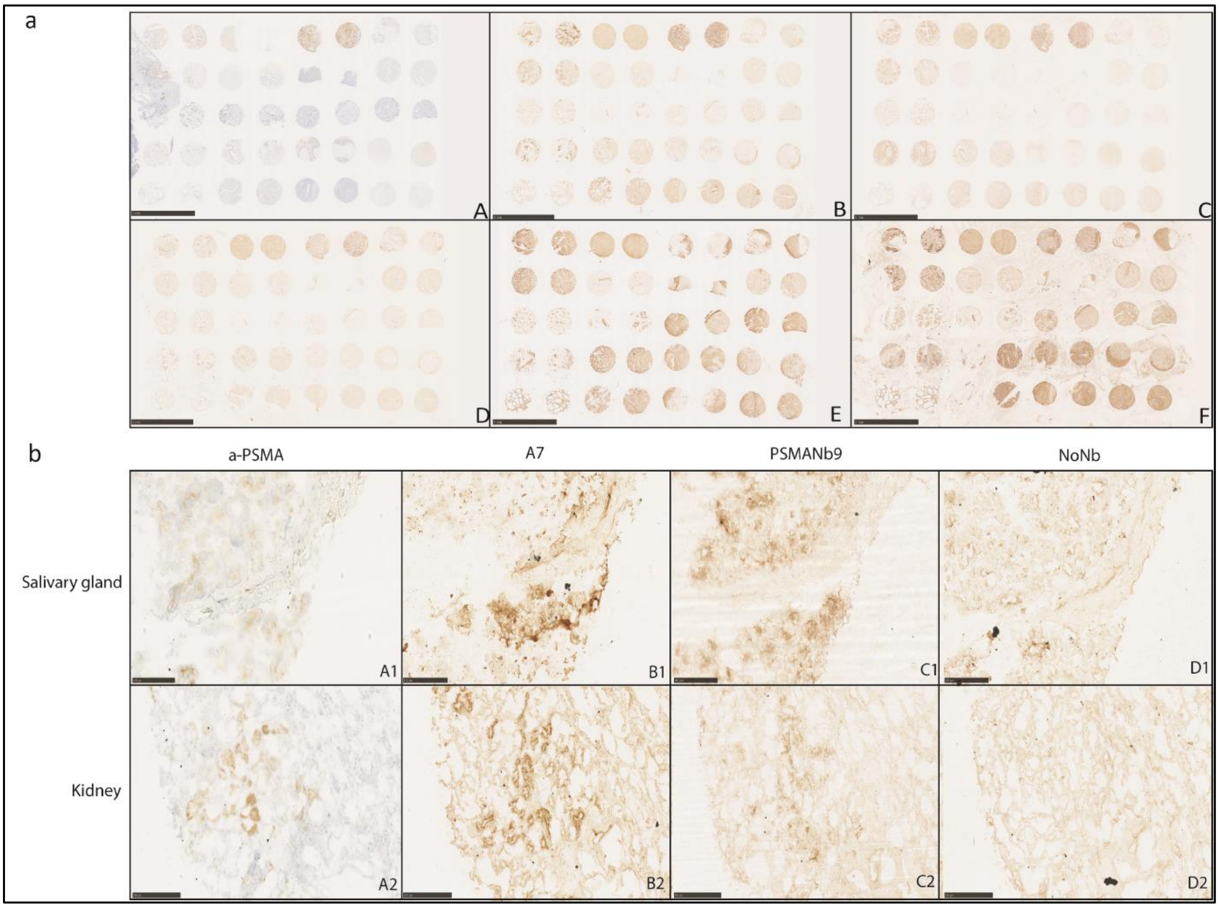


C

| | 1 | 2 | 3 | 4 | 5 | 6 | 7 | 8 |
|---|---|---|---|---|---|---|---|---|
| **A** | Salivary gland | Salivary gland | Liver | Liver | Small intestine | Small intestine | Stomach | Stomach |
| **B** | Kidney | Kidney | Skeletal Muscle | Skeletal Muscle | Skin | Skin | Heart | Heart |
| **C** | Placenta | Placenta | Breast | Breast | Cervix | Cervix | Uterus | Uterus |
| **D** | Spleen | Spleen | Lung | Lung | Brain, Cerebellum | Brain, Cerebellum | Brain, Cerebrum | Brain, Cerebrum |
| **E** | Thyroid | Thyroid | Pancreas | Pancreas | Ovary | Ovary | Adrenal | Adrenal |

*Suppl. Figure 4: **(A)** IHC staining of human normal tissue microarrays. A. anti-PSMA antibody. B. A7 nanobody-phage. C. PSMANb9 nanobody-phage. D. The phage without a nanobody (NoNb). E. anti-CD9 antibody. F. H6 nanobody-phage specific to CD9. Scale bar is 5 millimeters. **(B)** Enlarged sections of salivary gland (A1-D1) and kidney (A2-D2) tissues. Scale bars are 100 µm and 250 µm, respectively. All images were made by Nanozoomer 2.0 HT digital slide scanner (Hamamatsu Photonics K.K, Germany), visualized and analyzed using NDP viewer software. **(C)** Location of tissues on the TMA.*

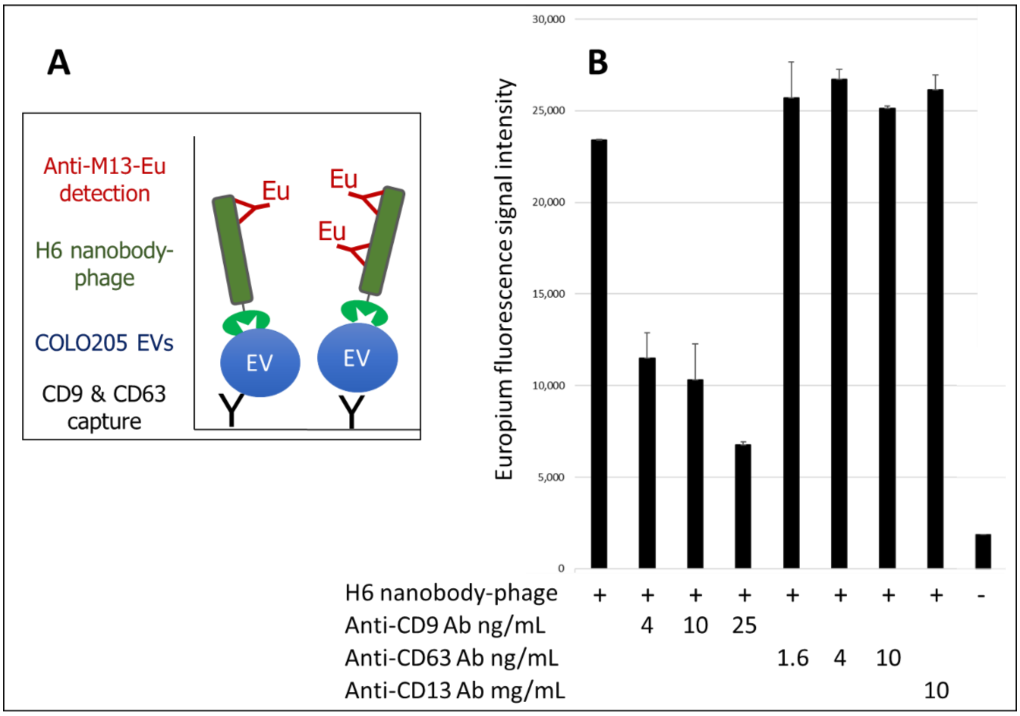


*Suppl. Figure 5: Time-Resolved Fluorescence ImmunoAssay (TR-FIA) demonstrating competition between the H6 nanobody-phage and an anti-CD9 antibody.* ***(A)*** *TR-FIA using anti-CD9 and anti-CD63 antibodies to capture COLO205 extracellular vesicles (EVs). Binding of the H6 nanobody-phage is measured using an Eu-conjugated anti-M13 antibody.* ***(B)*** *TR-FIA H6 nanobody-phage binding competition using different concentrations of antibodies against the tetraspanins CD9, CD63 and the aminopeptidase N (CD13). Representative experiment performed in duplicate (± SD).*

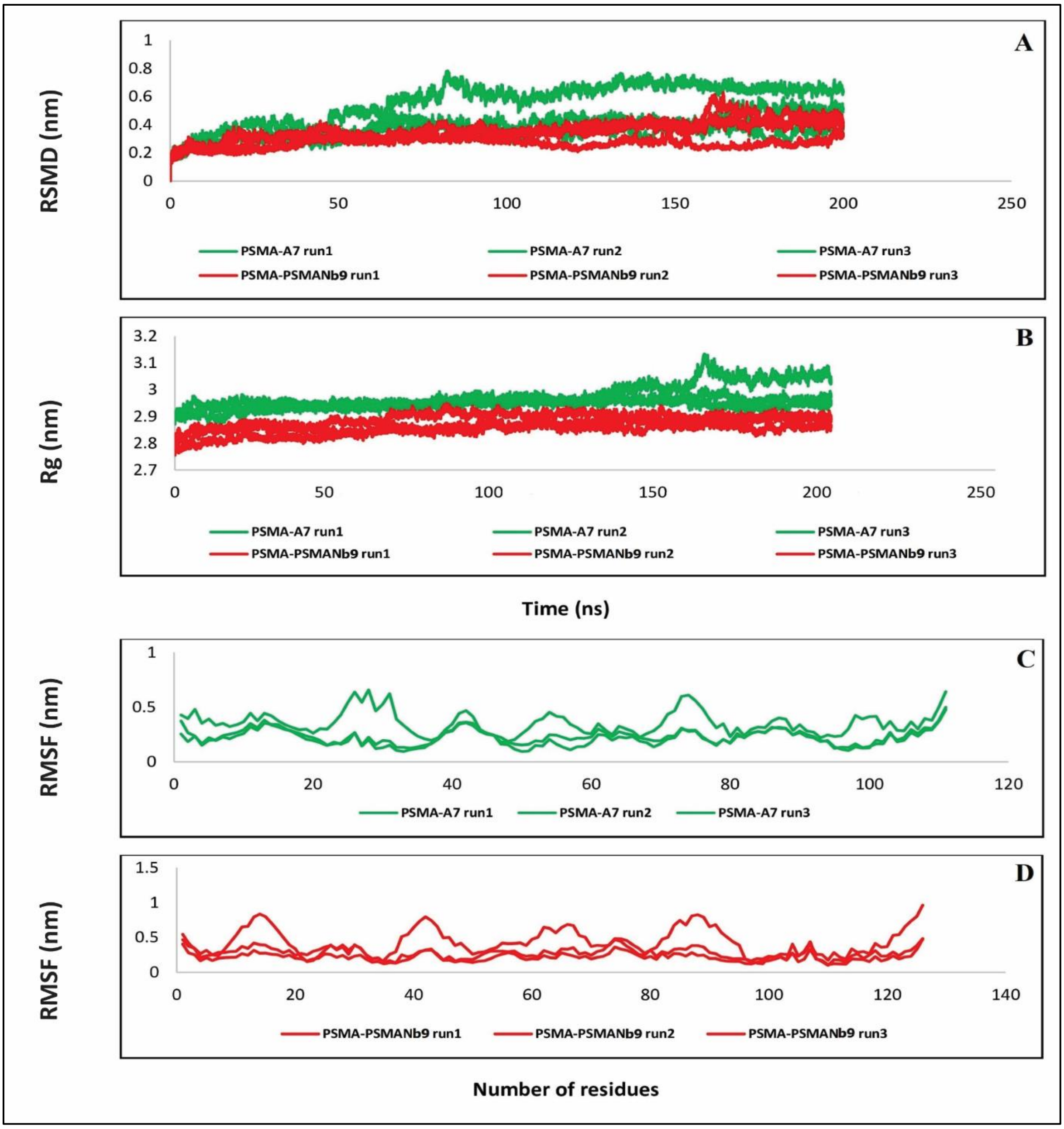


*Suppl. Figure 6: Evaluation of nanobody-PSMA complex stability during MD simulation. PSMANb9 and A7 are represented in red and green, respectively. (**A**) Calculated RMSD per unit time over the 200 ns simulation with three repetitions. Stability of both PSMANb9-PSMA and A7-PSMA complexes throughout the simulation. (**B**) Calculated Rg per unit time over the 200 ns simulation. (**C**) Backbone RMSF calculated over the 200 ns simulation for each residue of A7-PSMA complex. (**D**) Backbone RMSF calculated over the 200 ns simulation for each residue of PSMANb9-PSMA complex. Root Mean Square Deviation (RMSD), Radius of Gyration (Rg) and Root Mean Square Fluctuation (RMSF).*

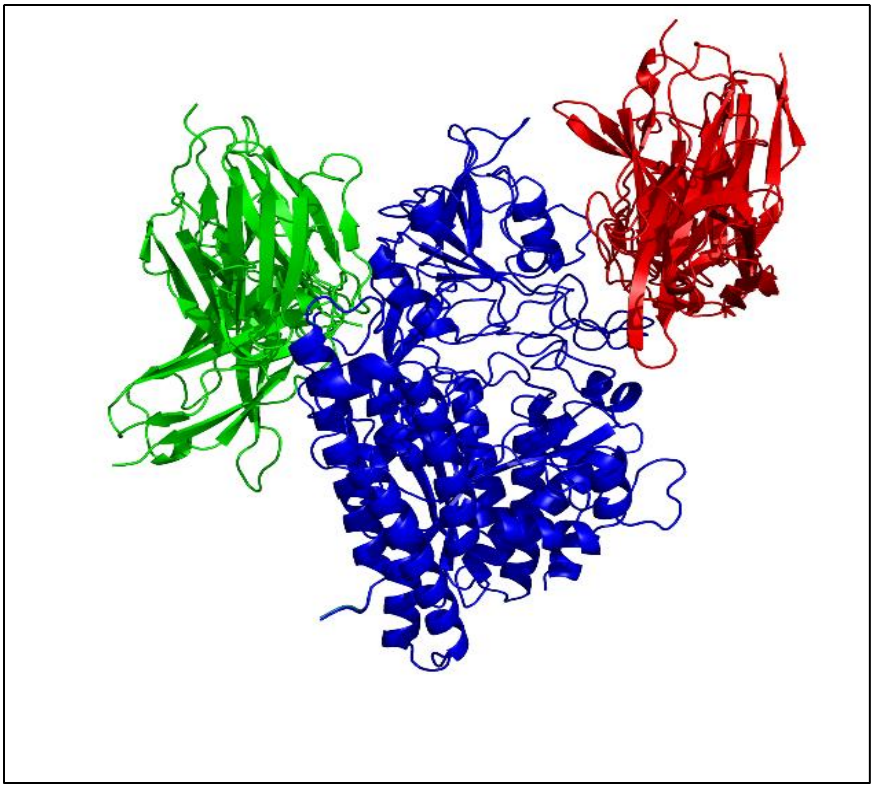

*Suppl. Figure 7. Docking models of nanobody-PSMA complexes. The previous docking of 6 nanobodies (Figure 5) identified 2 different epitopes. An additional 21 nanobodies with different CDR3 sequences were selected for folding and docking to identify potential novel epitopes on PSMA. All nanobodies tested so far, are predicted to bind to one of the epitopes bound by A7 or PSMANb9.*
*Example: Left part (green): A7, PSMANb12, 19. Right part (red): PSMANb9, 15, 21*

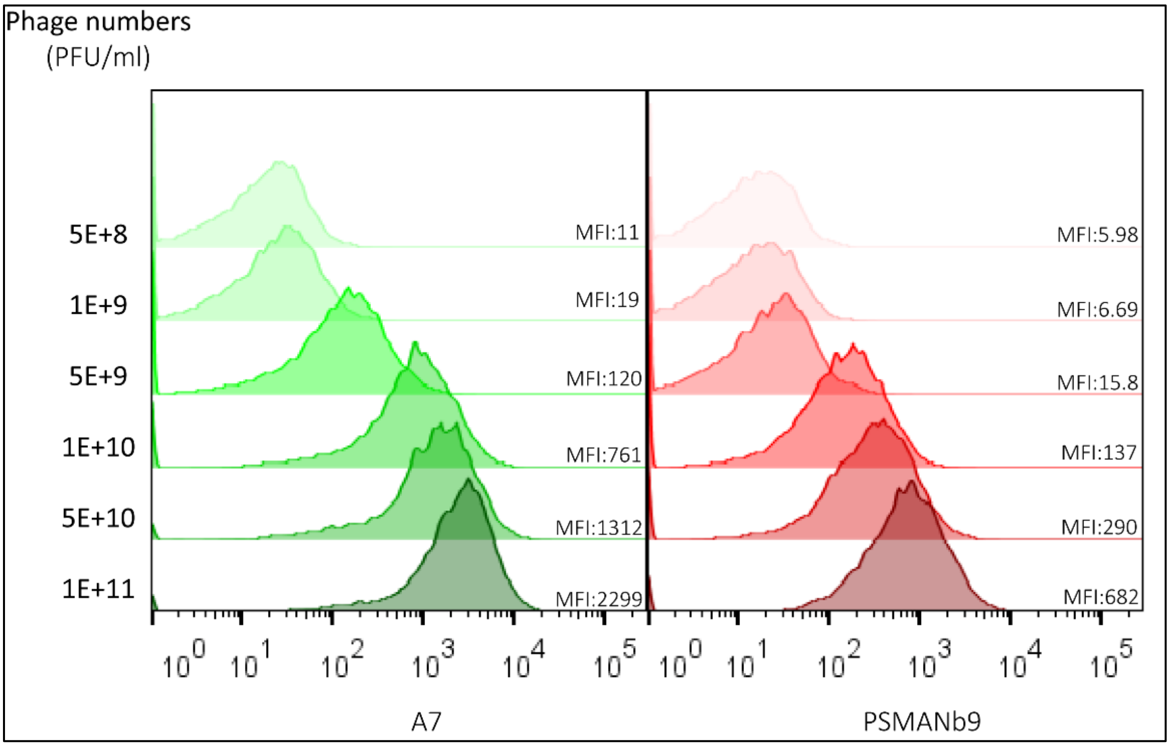


*Suppl. Figure 8: Flow cytometry to determine optimal concentration of ATTO488-conjugated A7 and PSMANb9 nanobody-phages. Per condition, 500,000 LNCaP cells were incubated for 1 hour at room temperature with different concentrations of ATTO488-labelled A7-phages (green) or PSMANb9-phages (red). The lowest concentration at which a clear shift in mean fluorescent intensity (MFI) and number of positive cells was observed, was selected for the competition experiments: 5E+9 for A7-phages and 1E+10 for the PSMANb9-phages.*

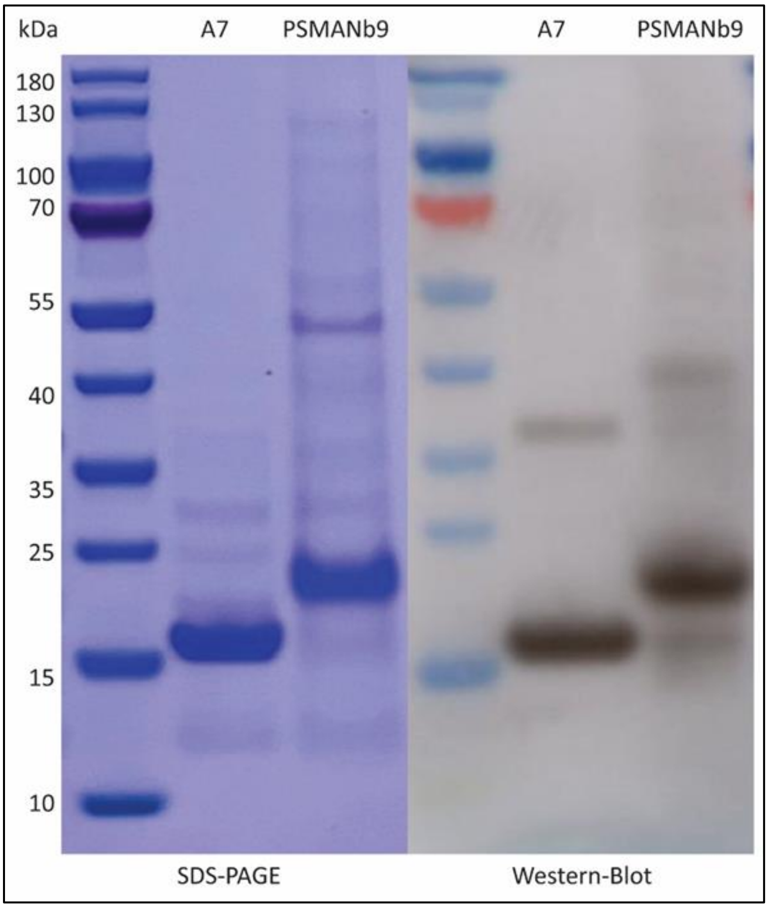


*Suppl. Figure 9: SDS-PAGE Coomassie blue staining (left) of bacterial produced A7 (~14.7 kDa) and PSMANb9 (~16.5 kDa) purified nanobody proteins. Protein bands on Western blot (right) were detected using anti-His tag antibody.*

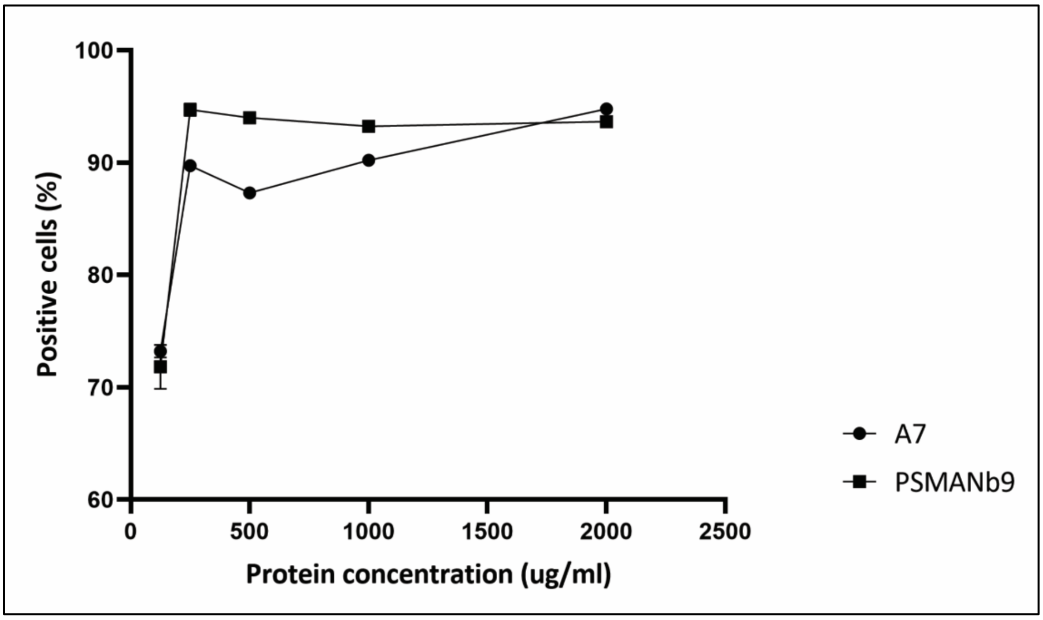


*Suppl. Figure 10: Epitope saturation using a series of purified A7 and PSMANb9 nanobody protein concentrations on LNCaP cell line. To determine the concentration of A7 and PSMANb9 proteins for optimal competition, flow cytometry was performed using 500,000 LNCaP cells incubated with different concentrations of nanobody proteins. The nanobodies bound to LNCaP cells were stained with anti-Myc tag antibody followed by goat-anti-mouse-AF647. The number of positive cells plateaued above 250 ug/mL. the concentration of 1 mg/mL was used for the competition experiments.*

#### 2. Supplementary Tables

| NAME | Library | LNCaP | PC346C | MDAPCa2b | 22Rv1 | MSKPCa12 | VCaP | PC339C | Hep3B | JON | T24 | MCF7 | T47D | HT29 | RKO | SW620 | HUVEC | SK-LMS-1 | SK-UT-1 | ST88-14 | 90-8TL | MLS1765-9 | MLS402-9 | OV90 | SKOV3 | BxPC-3 | PANC-1 | G-361 | HCC1806 | COLO205 | A549 | NCI-H1299 | NCI-H460 | NCI-H820 | MG-63 | SaOS-2 | U2OS | DU145 | PC3 | PNT2C2 | RWPE2 | 5637 | J82 | RT-112 | TCCSUP | B16-WT | B16-PSMA |
|---|---|---|---|---|---|---|---|---|---|---|---|---|---|---|---|---|---|---|---|---|---|---|---|---|---|---|---|---|---|---|---|---|---|---|---|---|---|---|---|---|---|---|---|---|---|---|---|
| | 1525 | 4929 | 3996 | 1075 | 2396 | 1032 | 352 | 324 | 1 | 1 | 1 | 1 | 429 | 1 | 1 | 1 | 1 | 698 | 1 | 260 | 178 | 1 | 1 | 1 | 1 | 1 | 369 | 119 | 630 | 445 | 294 | 1 | 968 | 122 | 525 | 1096 | 580 | 330 | 1 | 220 | 276 | 99 | 578 | 259 | 406 | 333 | 31496 |
| PSMANb1_L1P4 | 1 | 7068 | 7437 | 1935 | 2355 | 1032 | 69 | 1 | 1 | 1 | 1 | 1 | 1 | 1 | 1 | 1 | 1 | 1 | 1 | 1 | 1 | 1 | 1 | 1 | 1 | 1 | 1 | 1 | 1 | 89 | 1 | 1 | 1 | 122 | 1 | 274 | 1 | 1 | 1 | 110 | 1 | 1 | 1 | 86 | 1 | 1 | 33170 |
| | 180 | 7281 | 4141 | 5476 | | | 273 | | | 1 | 1 | 1 | 115 | | | | | | | | | | | | | | | | | | | | | | | | | | | | | 1 | 116 | 86 | 101 | 1 | 55235 |
| PSMANb6_LUPCa2 | 1 | 10922 | 7963 | 10570 | | | 273 | | | 97 | 1 | 1 | 1 | | | | | | | | | | | | | | | | | | | | | | | | | | | | | 1 | 116 | 1 | 1 | 1 | 110573 |
| | 549 | 5208 | 4440 | 860 | 1410 | 516 | 119 | 81 | 1 | 1 | 1 | 1 | 1 | 408 | 1 | 1 | 1 | 1 | 1 | 1 | 178 | 1 | 1 | 1 | 1 | 1 | 1 | 1 | 126 | 1 | 147 | 1 | 242 | 122 | 1 | 1 | 232 | 1 | 1 | 110 | 69 | 197 | 231 | 172 | 203 | 111 | 36270 |
| | 1 | 1765 | 1062 | 3693 | | | 91 | | | 1 | 1 | 1 | 1 | | | | | | | | | | | | | | | | | | | | | | | | | | | | | 1 | 1 | 1 | 1 | 1 | 19404 |
| | 1 | 1434 | 425 | 2420 | | | 1 | | | 1 | 1 | 1 | 1 | | | | | | | | | | | | | | | | | | | | | | | | | | | | | 1 | 1 | 1 | 1 | 1 | 13449 |
| | 1 | 1765 | 637 | 2547 | | | 1 | | | 1 | 1 | 1 | 1 | | | | | | | | | | | | | | | | | | | | | | | | | | | | | 1 | 116 | 1 | 1 | 1 | 12833 |
| | 1 | 1655 | 743 | 2165 | | | 1 | | | 1 | 1 | 1 | 1 | | | | | | | | | | | | | | | | | | | | | | | | | | | | | 1 | 1 | 1 | 1 | 1 | 12423 |
| | 1 | 2206 | 1274 | 3184 | | | 1 | | | 97 | 1 | 1 | 1 | | | | | | | | | | | | | | | | | | | | | | | | | | | | | 1 | 1 | 1 | 1 | 1 | 18993 |
| | 1 | 883 | 743 | 2674 | | | 91 | | | 1 | 1 | 1 | 1 | | | | | | | | | | | | | | | | | | | | | | | | | | | | | 1 | 1 | 1 | 101 | 172 | 15400 |
| PSMANb11_LUPCa1 | 18900 | 649023 | 244942 | 941744 | | | 6923 | | | 1462 | 1139 | 2437 | 3441 | | | | | | | | | | | | | | | | | | | | | | | | | | | | | 3454 | 3699 | 1552 | 2232 | 5254 | 4667055 |
| | 1 | 1545 | 531 | 2802 | | | 1 | | | 1 | 1 | 1 | 1 | | | | | | | | | | | | | | | | | | | | | | | | | | | | | 1 | 116 | 1 | 1 | 1 | 7289 |
| | 1 | 2537 | 1274 | 4585 | | | 91 | | | 1 | 1 | 1 | 1 | | | | | | | | | | | | | | | | | | | | | | | | | | | | | 1 | 116 | 1 | 1 | 1 | 18172 |
| | 1 | 1765 | 956 | 2038 | | | 1 | | | 1 | 1 | 1 | 1 | | | | | | | | | | | | | | | | | | | | | | | | | | | | | 1 | 1 | 1 | 1 | 1 | 12833 |
| | 610 | 8742 | 4329 | 2795 | 906 | 86 | 69 | 1 | 1 | 1 | 1 | 1 | 1 | 204 | 1 | 1 | 1 | 1 | 1 | 1 | 1 | 1 | 1 | 1 | 1 | 1 | 123 | 119 | 1 | 178 | 1 | 127 | 1 | 488 | 1 | 137 | 1 | 220 | 136 | 55 | 69 | 1 | 116 | 1 | 101 | 1 | 43090 |
| | 427 | 3720 | 2220 | 430 | 1662 | 172 | 103 | 1 | 1 | 1 | 1 | 357 | 1 | 204 | 1 | 1 | 243 | 1 | 1 | 260 | 178 | 1 | 426 | 1 | 125 | 1 | 1 | 1 | 378 | 89 | 147 | 1 | 121 | 1 | 210 | 137 | 116 | 1 | 1 | 55 | 69 | 1 | 231 | 1 | 1 | 222 | 21886 |
| | 366 | 5952 | 8547 | 1935 | 2760 | 258 | 46 | 81 | 1 | 1 | 1 | 1 | 1 | 1 | 1 | 1 | 1 | 1 | 1 | 1 | 1 | 1 | 1 | 1 | 1 | 1 | 123 | 1 | 1 | 1 | 1 | 1 | 121 | 1 | 1 | 1 | 1 | 1 | 1 | 55 | 138 | 1 | 1 | 172 | 1 | 111 | 44392 |
| | 305 | 6045 | 4551 | 2150 | 3396 | 516 | 78 | 81 | 1 | 1 | 1 | 1 | 1 | 1 | 1 | 1 | 1 | 349 | 1 | 1 | 1 | 1 | 1 | 176 | 125 | 1 | 1 | 1 | 126 | 267 | 147 | 1 | 121 | 1 | 105 | 137 | 348 | 1 | 1 | 55 | 207 | 99 | 1 | 172 | 1 | 444 | 47058 |
| PSMANb5_L1P4 | 854 | 9300 | 8436 | 1505 | 4719 | 258 | 69 | 81 | 401 | 1 | 1 | 1 | 1 | 1 | 1 | 1 | 243 | 1 | 1 | 260 | 178 | 1 | 1 | 1 | 1 | 1 | 1 | 238 | 1 | 178 | 1 | 254 | 121 | 1 | 1 | 1 | 116 | 1 | 272 | 55 | 276 | 197 | 1 | 1 | 1 | 222 | 67394 |
| | 305 | 5673 | 5883 | 860 | 2924 | 344 | 1 | 1 | 1 | 1 | 1 | 1 | 429 | 1 | 1 | 1 | 1 | 1 | 1 | 1 | 178 | 1 | 213 | 1 | 1 | 1 | 123 | 1 | 1 | 89 | 1 | 127 | 1 | 122 | 105 | 1 | 1 | 1 | 1 | 110 | 1 | 1 | 1 | 1 | 1 | 1 | 41726 |
| | 1 | 4192 | 1911 | 3948 | | | 1 | | | 1 | 1 | 1 | 1 | | | | | | | | | | | | | | | | | | | | | | | | | | | | | 99 | 1 | 172 | 1 | 1 | 21355 |
| | 549 | 16740 | 5439 | 1935 | 2587 | 774 | 182 | 1 | 401 | 1 | 415 | 1 | 1 | 1 | 1 | 1 | 1 | 1 | 1 | 1 | 1 | 1 | 1 | 1 | 1 | 185 | 1 | 1 | 252 | 356 | 1 | 1 | 1 | 244 | 105 | 274 | 348 | 110 | 1 | 110 | 69 | 99 | 1 | 86 | 101 | 111 | 57288 |
| PSMANb7_L1P4 | 6466 | 92814 | 49284 | 17630 | 19130 | 5848 | 1481 | 1296 | 401 | 814 | 415 | 357 | 858 | 1020 | 1 | 498 | 1458 | 2443 | 1182 | 260 | 534 | 864 | 852 | 880 | 750 | 925 | 1353 | 1071 | 2646 | 1068 | 1911 | 1524 | 1694 | 1220 | 1890 | 1370 | 812 | 1100 | 1632 | 1375 | 1518 | 296 | 809 | 690 | 710 | 1443 | 421290 |
| | 1 | 4082 | 1062 | 4203 | | | 91 | | | 1 | 1 | 1 | 1 | | | | | | | | | | | | | | | | | | | | | | | | | | | | | 99 | 1 | 172 | 1 | 1 | 23203 |
| | 61 | 5301 | 2109 | 1505 | 1823 | 86 | 1 | 1 | 1 | 1 | 1 | 1 | 1 | 1 | 1 | 1 | 1 | 1 | 1 | 1 | 1 | 1 | 1 | 1 | 1 | 1 | 1 | 1 | 1 | 1 | 1 | 1 | 1 | 1 | 1 | 1 | 1 | 1 | 1 | 1 | 1 | 1 | 1 | 1 | 1 | 1 | 8866 |
| | 305 | 26877 | 14097 | 3440 | 8055 | 3268 | 258 | 1 | 1 | 1 | 1 | 1 | 1 | 1 | 1 | 1 | 243 | 1 | 1 | 1 | 1 | 1 | 1 | 1 | 1 | 1 | 1 | 1 | 504 | 1 | 147 | 127 | 1 | 122 | 210 | 274 | 116 | 330 | 1 | 55 | 138 | 1 | 1 | 1 | 1 | 111 | 95108 |
| | 183 | 6231 | 4884 | 430 | 1626 | 86 | 1 | 162 | 1 | 1 | 1 | 1 | 1 | 1 | 1 | 1 | 1 | 349 | 1 | 1 | 1 | 1 | 1 | 1 | 1 | 1 | 1 | 119 | 1 | 1 | 1 | 1 | 1 | 122 | 1 | 1 | 1 | 1 | 136 | 55 | 1 | 1 | 1 | 1 | 1 | 1 | 32302 |
| PSMANb2_L1P4 | 610 | 29388 | 19425 | 6020 | 8211 | 774 | 179 | 1 | 401 | 1 | 1 | 1 | 1 | 1 | 1 | 1 | 1 | 698 | 1 | 1 | 1 | 1 | 1 | 1 | 250 | 1 | 1 | 119 | 1 | 267 | 1 | 254 | 1 | 1 | 210 | 137 | 1 | 220 | 136 | 55 | 276 | 99 | 116 | 1 | 1 | 1 | 129022 |
| PSMANb9_LUPCa1 | 1 | 3751 | 2761 | 6622 | | | 182 | | | 1 | 1 | 1 | 1 | | | | | | | | | | | | | | | | | | | | | | | | | | | | | 1 | 116 | 1 | 101 | 1 | 50410 |
| A7_L1P4 | 549 | 34131 | 18093 | 11180 | 10901 | 3010 | 273 | 81 | 1 | 1 | 1 | 1 | 1 | 204 | 1 | 1 | 1 | 1 | 591 | 260 | 1 | 1 | 1 | 1 | 1 | 370 | 1 | 119 | 1 | 1 | 1 | 127 | 363 | 1 | 105 | 1 | 1 | 110 | 1 | 1 | 207 | 1 | 116 | 1 | 1 | 555 | 151404 |
| | 854 | 15624 | 10878 | 7525 | 4812 | 774 | 23 | 81 | 1 | 1 | 1 | 1 | 1 | 204 | 1 | 1 | 486 | 1 | 1 | 260 | 1 | 432 | 1 | 1 | 1 | 1 | 1 | 1 | 126 | 89 | 1 | 1 | 1 | 366 | 420 | 1 | 1 | 1 | 136 | 1 | 138 | | | | | 333 | 54002 |
| | 1952 | 7068 | 4329 | 1290 | 2082 | 946 | 205 | 324 | 1 | 407 | 415 | 357 | 429 | 1 | 378 | 1 | 243 | 1 | 1 | 1 | 178 | 1 | 213 | 352 | 125 | 1 | 123 | 238 | 378 | 445 | 588 | 254 | 363 | 488 | 840 | 411 | 464 | 1 | 408 | 110 | 414 | 395 | 462 | 172 | 203 | 222 | 58528 |
| | 3233 | 3999 | 2553 | 5375 | 1781 | 258 | 478 | 486 | 1 | 1 | 415 | 357 | 429 | 408 | 1 | 1 | 486 | 1 | 1 | 260 | 356 | 864 | 213 | 880 | 250 | 925 | 615 | 1071 | 756 | 712 | 1176 | 381 | 121 | 732 | 525 | 274 | 580 | 220 | 1 | 330 | 759 | 197 | 1 | 259 | 203 | 1776 | 29016 |
| | 244 | 3999 | 3219 | 2150 | 1679 | 344 | 55 | 1 | 1 | 1 | 1 | 1 | 1 | 1 | 1 | 1 | 1 | 1 | 1 | 1 | 1 | 1 | 1 | 1 | 1 | 1 | 1 | 1 | 1 | 1 | 1 | 1 | 1 | 1 | 105 | 274 | 1 | 1 | 1 | 55 | 1 | 1 | 1 | 1 | 1 | 444 | 28830 |
| A5_L1P4 | 488 | 10137 | 6105 | 1720 | 885 | 86 | 23 | 1 | 802 | 1 | 1 | 1 | 1 | 1 | 1 | 1 | 1 | 1 | 1 | 1 | 178 | 432 | 213 | 1 | 1 | 185 | 1 | 1 | 126 | 1 | 1 | 1 | 1 | 366 | 210 | 1 | 1 | 110 | 136 | 1 | 69 | 99 | 116 | 86 | 101 | 1 | 64790 |
| | 220 | 1986 | 425 | 2038 | | | 1 | | | 1 | 1 | 1 | 1 | | | | | | | | | | | | | | | | | | | | | | | | | | | | | 1 | 231 | 1 | 1 | 1 | 10575 |
| | 330 | 4854 | 743 | 3693 | | | 1 | | | 1 | 1 | 1 | 1 | | | | | | | | | | | | | | | | | | | | | | | | | | | | | 99 | 1 | 1 | 1 | 1 | 29158 |
| | 330 | 4744 | 1486 | 4203 | | | 91 | | | 1 | 1 | 1 | 1 | | | | | | | | | | | | | | | | | | | | | | | | | | | | | 99 | 1 | 86 | 1 | 1 | 24640 |
| | 330 | 7943 | 1274 | 8660 | | | 91 | | | 1 | 1 | 143 | 1 | | | | | | | | | | | | | | | | | | | | | | | | | | | | | 1 | 1 | 86 | 101 | 86 | 36139 |
| PSMANb10_LUPCa1 | 23020 | 135696 | 20598 | 110921 | | | 2277 | | | 1364 | 1449 | 1577 | 1721 | | | | | | | | | | | | | | | | | | | | | | | | | | | | | 1579 | 1040 | 1207 | 1420 | 1981 | 603274 |
| | 1708 | 22320 | 16983 | 78045 | 89704 | 4128 | 5547 | 4779 | 6015 | 1628 | 1 | 16065 | 1716 | 612 | 1890 | 2490 | 243 | 698 | 1182 | 2080 | 534 | 1296 | 852 | 7568 | 2250 | 9065 | 738 | 2499 | 1764 | 4539 | 1617 | 635 | 726 | 366 | 840 | 1370 | 26912 | 30580 | 70448 | 94050 | 621 | 296 | 578 | 259 | 406 | 444 | 88722 |
| | 549 | 5766 | 2886 | 7095 | 1215 | 1 | 80 | 1 | 1 | 1 | 1 | 1 | 1 | 204 | 1 | 1 | 243 | 1 | 591 | 1 | 1 | 1 | 1 | 1 | 1 | 1 | 123 | 1 | 1 | 1 | 1 | 127 | 121 | 122 | 105 | 1 | 116 | 110 | 1 | 1 | 69 | 1 | 231 | 1 | 1 | 111 | 42284 |
| | 2684 | 6603 | 2442 | 12470 | 3081 | 1032 | 1807 | 567 | 1 | 1 | 1 | 1785 | 1287 | 204 | 378 | 1494 | 2187 | 698 | 1 | 260 | 178 | 864 | 426 | 352 | 2750 | 1 | 738 | 1071 | 1386 | 712 | 1029 | 1 | 1089 | 732 | 1470 | 959 | 580 | 550 | 136 | 220 | 1794 | 296 | 347 | 690 | 101 | 111 | 41912 |
| | 1037 | 11253 | 2775 | 12685 | 2775 | 2580 | 321 | 243 | 1 | 407 | 1 | 1 | 1 | 1 | 1 | 1 | 243 | 1 | 1 | 1 | 1 | 1 | 1 | 1 | 1 | 185 | 246 | 357 | 252 | 267 | 1 | 1 | 121 | 244 | 105 | 1 | 1 | 440 | 1 | 165 | 345 | 99 | 116 | 86 | 1 | 444 | 67580 |
| | 244 | 5673 | 1776 | 7955 | 2393 | 258 | 280 | 243 | 401 | 1 | 1 | 1 | 1 | 1 | 1 | 498 | 1 | 1 | 591 | 260 | 1 | 1 | 213 | 1 | 1 | 185 | 123 | 1 | 378 | 89 | 1 | 254 | 121 | 122 | 105 | 274 | 348 | 1 | 1 | 1 | 69 | 1 | 1 | 86 | 304 | 333 | 54188 |
| | 427 | 3348 | 777 | 6665 | 803 | 258 | 78 | 81 | 1 | 1 | 1 | 1 | 429 | 1 | 1 | 1 | 1 | 1 | 1 | 1 | 1 | 432 | 213 | 176 | 1 | 1 | 123 | 1 | 1 | 1 | 147 | 254 | 121 | 1 | 105 | 1 | 116 | 1 | 136 | 1 | 1 | 1 | 231 | 172 | 101 | 222 | 23684 |
| | 610 | 2976 | 777 | 4085 | 906 | 172 | 250 | 1 | 401 | 1 | 1 | 1 | 1 | 1 | 1 | 1 | 243 | 349 | 1 | 1 | 356 | 1 | 1 | 528 | 250 | 1 | 1 | 119 | 1 | 1 | 1 | 127 | 1 | 1 | 210 | 137 | 116 | 1 | 136 | 165 | 69 | 99 | 1 | 86 | 101 | 111 | 16058 |
| | 122 | 1302 | 777 | 3440 | 1439 | 172 | 818 | 81 | 1 | 1 | 1 | 1 | 1 | 1 | 1 | 1 | 1 | 1 | 1 | 1 | 1 | 1 | 1 | 1 | 1 | 1 | 1 | 119 | 1 | 178 | 1 | 1 | 1 | 1 | 1 | 137 | 1 | 220 | 1 | 55 | 69 | 1 | 1 | 1 | 1 | 111 | 1240 |
| PSMANb3_L1P4 | 427 | 4743 | 1443 | 5805 | 1384 | 86 | 170 | 1 | 1 | 1 | 1 | 1 | 1 | 204 | 378 | 1 | 1 | 349 | 1 | 260 | 356 | 432 | 213 | 176 | 125 | 1 | 123 | 119 | 126 | 1 | 147 | 127 | 121 | 1 | 105 | 137 | 116 | 110 | 1 | 55 | 1 | 1 | 1 | 86 | 101 | 111 | 33542 |
| | 1037 | 19251 | 7881 | 26230 | 5710 | 430 | 1054 | 81 | 1 | 1221 | 1 | 357 | 429 | 1 | 1 | 1 | 243 | 1 | 1 | 1 | 178 | 432 | 1 | 352 | 375 | 1110 | 123 | 238 | 378 | 178 | 294 | 254 | 1 | 1 | 1 | 274 | 116 | 220 | 816 | 495 | 1794 | 888 | 116 | 172 | 101 | 2109 | 144646 |
| | 23058 | 107043 | 24087 | 27735 | 22981 | 7052 | 5819 | 3888 | 6817 | 3256 | 7055 | 4284 | 6435 | 4284 | 5670 | 6972 | 5346 | 4886 | 4728 | 7020 | 5518 | 6912 | 5325 | 4048 | 4750 | 3515 | 2337 | 2975 | 5040 | 5251 | 4263 | 4699 | 4719 | 3416 | 8400 | 6987 | 4640 | 5170 | 5304 | 4235 | 4761 | 1579 | 1734 | 1466 | 1420 | 19203 | 273730 |

*Suppl. Table 1: Top 52 nanobody selected against PSMA and binding data in cell lines, the L1P4 library and the B16-WT and B16-PSMA 1 round single cell panning.*
*Provided as an Excel file.*

| Nanobody | 3D modeling tool | ERRAT score | MolProbity score | PROCHECK- Residues in allowed region (%) | ProSA-web Z-Score |
|---|---|---|---|---|---|
| A3 | Robetta | 88.5 | 0.50 | 92 | -6.33 |
| | SWISS-MODEL | 78.1 | 0.68 | 97 | -6.6 |
| A7 | Robetta | 100 | 0.76 | 88 | -6.5 |
| | SWISS-MODEL | 100 | 1.19 | 96 | -6.13 |
| PSMANb5 | Robetta | 82.5 | 0.68 | 92.4 | -6.85 |
| | SWISS-MODEL | 86.1 | 0.91 | 96 | -6.7 |
| PSMANb6 | Robetta | 92 | 1.06 | 93.1 | -7.35 |
| | SWISS-MODEL | 89.7 | 1.35 | 93 | -7.15 |
| PSMANb9 | Robetta | 85.3 | 0.81 | 94 | -6.72 |
| | Swiss Model | 74.1 | 0.68 | 94 | -5.17 |
| PSMANb11 | Robetta | 87.2 | 0.92 | 90.3 | -7.35 |
| | SWISS-MODEL | 90.5 | 0.82 | 94.2 | -6.03 |

*Suppl. Table 2: Evaluation of the overall quality of nanobody models generated by Robetta and SWISS-MODEL. All models were considered as reliable and high quality, irrespective of the modeling tool. ERRAT, MolProbity and ProSA-web Z-score for all models suggested high structural quality, comparable to crystallographic structures. Analysis of Ramachandran plot by PROCHECK did not show a lot of residues out of the allowed region, further indicating the quality of models.*

| Nanobody | HADDOCK score | Docking pose of nanobodies in complex with PSMA (blue) |
|---|---|---|
| **A3** | -83 | 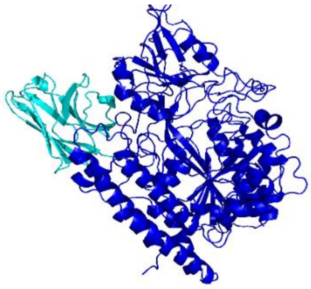 |
| **A7** | -120 | 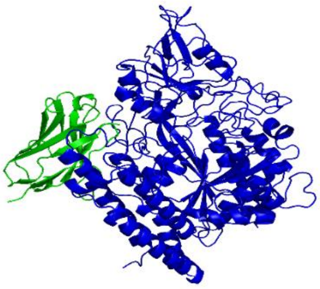 |

| | | |
|---|---|---|
| **PSMANb5** | -93 | 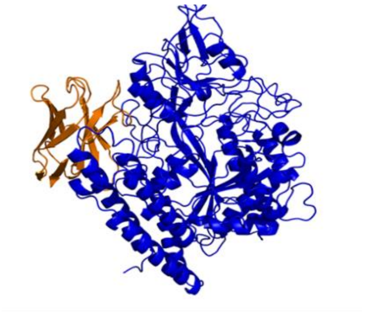 |
| **PSMANb6** | -93 | 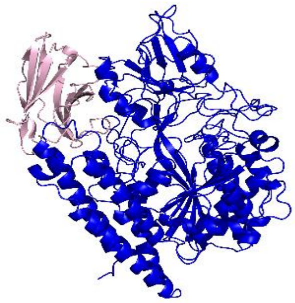 |
| **PSMANb9** | -117 | 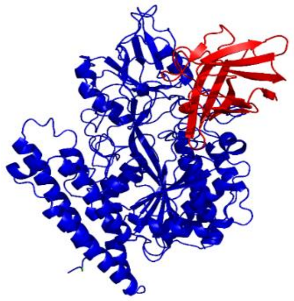 |
| **PSMANb11** | -90 | 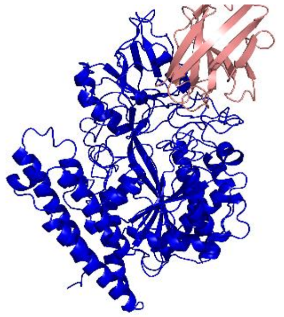 |

*Supp. Table 3: Docking analysis of nanobody-PSMA complexes: HADDOCK scores and structural models. The selected nanobodies could be clustered in two distinct groups, based on similar PSMA epitopes. More specifically, A3, A7, PSMANb5 and PSMANb6 shared a common epitope on PSMA, while PSMANb9 and PSMANb11 bind to a distinct epitope on the opposite side of PSMA.*

| Complex | HADDOCK score | Z-score | Cluster size | RMSD |
|---|---|---|---|---|
| **A3** | -83 | -1.4 | 19 | 0.4 |
| **A7** | -120 | -2.6 | 29 | 0.6 |
| **PSMANb5** | -93 | -1.5 | 20 | 0.9 |
| **PSMANb6** | -93 | -1.5 | 18 | 0.7 |
| **PSMANb9** | -117 | -1.9 | 53 | 0.6 |
| **PSMANb11** | -90 | -1.4 | 38 | 0.5 |

*Supp. Table 4: HADDOCK 2.4 scores and statistics of the top models for the nanobody-PSMA complexes. RMSD: Root Mean Square Deviation.*

| Antibodies and other detection molecules | Source | Identifier | Dilution | Application |
|---|---|---|---|---|
| Anti-M13 bacteriophage g8p antibody, mouse monoclonal [RL-ph2] | Abcam | ab9225 | 1:2000/1:200 | ELISA/IHC |
| Anti-PSMA antibody, mouse monoclonal | Sigma-Aldrich | SAB4200257 | 1:100 | Flow cytometry/IHC |
| Anti-PSMA antibody, rabbit monoclonal | Abcam | EP3253 | 1:5000 | Western Blot |
| Anti-CD9 antibody, mouse monoclonal (eBioSN4 (SN4 C3-3A2)) | Thermo Fisher | 14-0098-82 | 1:150/1:100 | ELISA/IHC |
| Streptavidin-biotin HRP | Dako | ab7403 | 1:1000 | ELISA/IHC |
| Goat-anti-mouse HRP antibody | Dako | P0447 | 1:2000/1:250/1:2000 | ELISA/IHC/Western blot |
| Goat-anti-rabbit HRP antibody | Dako | P0448 | 1:2000 | Western Blot |
| Goat-anti-mouse AF647 | Thermo Fisher | A32728 | 1:1000 | Flow cytometry |
| Anti-human GADPH antibody, mouse monoclonal (0411) | Santa Cruz Biotechnology | sc-47724 | 1:10000 | Western Blot |
| Anti Beta-actin antibody, mouse monoclonal (15G5A11/E2) | Thermo Fisher | MA1-140 | 1:10000 | Western Blot |
| Anti-myc-tag antibody, mouse monoclonal | Sigma-Aldrich | M4439 | 1:100/1:5000 | Flow cytometry/Western blot |

*Suppl. Table 5: Information on the antibodies used in this study.*

### 3. Supplementary Materials and Methods

**Llama immunization and VHH-library construction**

Three llamas were immunized to generate nanobodies targeting PCa antigens. One llama was immunized with four PCa cell lines (LNCaP, PC346C, VCaP, and MDAPCa2b) by Dr. Patrick Chames as previously described (L1P4 library) [1]. The other two llamas, LUPCa1 and LUPCa2, were immunized by Eurogentec, Belgium with cells isolated from a collection of fresh frozen PCa and bladder cancer (BlCa) patient tumors. Frozen tumor tissues were provided by the Erasmus MC Tissue Biobank in compliance with the Code for Proper Secondary Use of Human Tissue in the Netherlands and approved by the Erasmus MC Medical Ethics Committee according to the Medical Research Involving Human Subjects Act (METC-2004-261).

Blood was collected after the second or third immunization and peripheral blood lymphocytes were isolated. The LUPCa1 and LUPCa2 library constructions was performed by QVQ, The Netherlands. Briefly, RNA was isolated and cDNA was synthetized with reverse transcriptase. VHH (variable domains of camelid heavy-chain antibodies) genes were amplified by PCR introducing NotI and SfiI restriction enzyme sites (forward and reverse primers). VHH fragments were isolated from a 1% agarose gel, digested with SfiI and NotI, ligated into the pHEN1-6HisGS phagemid and transformed into E. *coli* TG1 cells to generate a library of 3.0E+9, 1.2E+9 and 0.6E+9 transformants for L1P4, LUPCa1 and LUPCa2, respectively.

**Cell culture**

Most of the cell lines in this study were collected from various research groups within the Erasmus MC and had different origins such as created by the research group (PC346C, PC339C), purchased from ATCC (American Type Culture Collection, USA; RWPE1, CRL-3607) and kind gifts from collaborators such as Dr. Marco Colombatti (University of Verona; B16-PSMA), Dr. Norman Maitland (University of York; PNT2C2), Dr. Nora Navone (MD Anderson Cancer Center; MDAPCa2b), Dr. Charles L Sawyers (Memorial Sloan Kettering Cancer Center; MSKPCa12D) and Dr. Ken Pienta (University of Michigan; VCaP).

Cell culture media and reagents were obtained from Gibco unless stated otherwise. LNCaP, 22Rv1, VCaP and DU145 cells were cultured in RPMI 1640 medium containing 10% (5% for DU145) fetal calf serum (FCS) and 1% penicillin-streptomycin (P/S). PC346C cells were cultured in Dulbecco's modified Eagle's F12 medium (DMEM), supplemented with 5% FCS, 0.01% BSA (Boehringer-Mannheim), 1% P/S, 1% insulin-transferrin-selenium (Life Technologies), 0.1 µg/mL fibronectin (Alfa Aesar), 0.1 nM R1881 androgen, 10 ng/mL epidermal growth factor, 0.5 µg/mL dexametason, 1 nM triiodothyronine, 0.1 mM phosphoethanolamine, 50 ng/mL cholera toxin and 20 µg/mL fetuin (all from Sigma). B16-WT and B16-PSMA cells were cultured in DMEM containing 10% FCS, 10 mM HEPES (Sigma), 20 µM 2-mercaptoethanol (Sigma) and 2 mM glutamine. For B16-PSMA, 1200 µg/mL geneticin (G418, Gibco) was added to this medium. MDA-PCa2b cells were grown in Ham's F12 medium containing 15% FCS, 1% P/S, 25 ng/mL cholera toxin, 10 ng/mL epidermal growth factor, 5 µM phosphoethanolamine, 120 pg/mL hydrocortisone and 1% insulin-transferrin-selenium. HEK293T cells were cultured in 1:1 ratio of DMEM and Ham's F12 medium, with 5% FCS and 1% P/S. Hep3B, T47D, MCF7, PANC-1, COLO205, BPH-1 and SK-LSM1 cells were maintained in DMEM supplemented with 10% FBS and 1% P/S. Bladder cancer cell lines (5637, RT112, T24, TCCSUP, J82) were cultured in RPMI 1640, supplemented with 10% FBS and 1% P/S. All cells were maintained at 37°C in a humidified atmosphere containing 5% $CO_2$. Cells

were detached and collected for panning, cell ELISA or IHC at 80% confluency using 3.4 mM EDTA in PBS.
Cell cultures were regularly tested for mycoplasma infection (MycoAlert Mycoplasma Detection kit, Lonza).

**Screening and selection of nanobodies using phage display technology**
All cell lines were blocked with PBS/10% FCS/2% BSA for 30 min at RT prior to selection, irrespective of the biopanning strategy. For 3R-NegPos, 1E+7 B16-WT cells were incubated with 2E+11 phages from the L1P4 library in PBS/10% FCS/2% BSA for 45 min at RT. B16-WT cells were then spun down, and the supernatant containing unbound phages was incubated with 1E+7 B16-PSMA cells in PBA/2% BSA for 45 min at RT. Cells were washed ten times with PBS/0.5% BSA and two times with PBS. Phages bound to PSMA-B16 cells were eluted with 200 mM triethylamine (Sigma) and reamplified in E. *coli* TG1 cells. Colonies were grown on 2YT (16 g/L tryptone, 10 g/L yeast extract and 5 g/L NaCl)/ampicillin (100 µg/mL)/glucose (2%) agar plates. Colonies were scraped from the 2YT plates, and stored at –80°C in the presence of 20% glycerol. The phage pool from first round of biopanning was titrated in E. *coli* TG1 cells and used to conduct the second round following the same negative-positive selection. Accordingly, the third biopanning round was performed using the phage pool form the second round.
For the 1R-SC, B16-WT and B16-PSMA cells were resuspended after detachment in PBS/2% BSA. Cell stocks of 5E+6 cells were prepared and stored at –80°C. For biopanning, cells were defrosted and blocked. Then 1E+7 B16-WT and B16-PSMA cells were incubated with 2E+11 phages, separately. Bound phages were eluted, amplified and stored following the same protocol as 3R-NegPos.
For the 1R-Cell Line Panning, The FOLH1 expression values of the cell line panel was retrieved from the CCLE RNAseq repository (22Q2; [2]) and from our own RNAseq data [3]. The cell lines with relative FOLH1 expression above log2 TPM of 5 were selected as PSMA expressing cell lines (PSMApos: VCaP, 22Rv1, LNCaP, MDAPCa2b, PC346C) while cell lines with lower expression than log2 TPM of 0.4 were selected as PSMA negative cell lines (PSMAneg: PC3, SaOS2, MG63, H460, H1299, A549, HT29, OV90, SK-UT-1, SK-LMS-1, SW620, RKO, MCF7, T24, PC339C). VCaP, 22Rv1, MDAPCa2b, PC346C and PC339 cells carry the Xenotropic Murine leukemia virus-Related Virus (XMRV) contamination. In order to eliminate selecting nanobodies against XMRV, the average nanobody binding level to XMRV-positive cell lines and the XMRV negative cell lines LNCaP, DU145 and PC3 were calculated and the ratio determined.
The B16-WT and B16-PSMA cell lines likely contain the MelARV [4] or Bxv-1 viruses (https://www.sigmaaldrich.com/deepweb/assets/sigmaaldrich/product/documents/406/200/scc420ds-ver-1-0.pdf).

For NGS, phagemids were isolated from the E. *coli* TG1 cells infected with the eluted nanobody-phages for each individual panning. To extract and amplify the nanobody cDNAs, PCR was performed using primers with the adapter, Illumina forward or reverse sequencing primer, a unique sample index combined with a heterogeneity spacer $(N)_{12\text{-}15}$ and the sequence binding the phagemid [5].
Forward primer:
5’ AATGATACGGCGACCACCGAGATCTACACTCTTTCCCTACACGACGCTCTTCCGATCT $(N)_{12\text{-}15}$ CTGGATTGTTATTACTCG 3’
Reverse primer:
5’ CAAGCAGAAGACGGCATACGAGATGTGACTGGAGTTCAGACGTGTGCTCTTCCGATCT $(N)_{12\text{-}15}$ GAGATGAGTTTTTGTTC 3’

PCR products were checked on gel for amplicon size and quantified using Bioanalyzer 2100 (Agilent) and pooled for Illumina MiSeq NGS.

**NGS processing pipeline**

The MiSeq fastq R1 and R2 files were first merged using PEAR (version 1.10.13) followed by demultiplexing and removal of adapter sequences (Cutadapt version 3.5). Sequences were aligned to the alpaca VHH reference using MiXCR (version 3.0.13). For each sample, nanobodies with the identical CDR3 amino acid sequence were grouped and counted, allowing a single nanobody in one cluster. The CDR3 clusters from each sample were merged into a table and corrected for differences in sequencing depth by normalizing cluster counts in each sample to the sum of 100 million reads in each sample. A custom R script was used to identify all same-length CDR3 clusters that are a single amino acid different (SaaDiff) to define CDR3 families with Levenshtein distance of 1.

**Nanobody-phage production**

The DNA sequence of the most prevalent full length nanobody was retrieved from sequencing data to be synthesized and inserted into pHEN1-6HISGS phagemid by Twist Bioscience (California, USA). The phagemid was transformed into chemically competent E. *coli* TG1 cells. Transformed colonies were grown in 2YT (16 g/L tryptone, 10 g/L yeast extract and 5 g/L NaCl)/ampicillin (100 µg/mL)/glucose (2%) overnight. The culture was used to inoculate 10 ml 2YT/ampicillin (100 µg/mL)/glucose (2%) until the optical density at 600 nm (OD600) nm reached 0.8. M13 K07ΔpIII hyperphages (Progen Biotechnik) were added with a Multiplicity of Infection (MOI) of 5 and the bacteria were pelleted and resuspended in TAK medium and grown overnight at 30°C. The bacteria were then pelleted and the supernatant containing the phage particles was mixed with 1/5 (V/V) of 20% W/V PEG and 2.5 M NaCl to precipitate the phages for 2 hrs on ice. Phages were pelleted at 4500xg for 30 min. The phage pellet was resuspended in PBS, filtered through a MF-Millipore Membrane filter 0.45 uM (Merck, US) and re-precipitated for 1 hr on ice. Finally, the phage pellet was resuspended in PBS and titrated using E. *coli* TG1 cells. The nanobody-phage stocks were stored at 4°C.

**Whole Cell ELISA**

The experiment was performed using two PSMA-positive cell lines, LNCaP and B16-PSMA, along with two PSMA-negative cell lines, DU145 and B16-WT. The cell ELISA protocol is described in the supplementary data. Initially, 50,000 cells were added in triplicates in a V-bottom 96-well plate (Nunc A/S, ThermoFisher Scientific, Denmark) in 2% BSA-PBS. 5E+10 nanobody-phages were then added to each well and incubated for 45 minutes at room temperature. Wells containing cells without phages added or with the phage without a nanobody (NoNb) added, served as negative control to measure the background signal. Mouse anti-PSMA antibody (Sigma) served as positive control. After washing 3 times with PBS-2% BSA, mouse anti-M13 antibody (Abcam, USA) was added and incubated for 45 minutes at room temperature. Binding of phages or antibody to the cells were detected by goat anti-mouse HRP antibody (Dako, Agilent, USA) followed by OPD substrate (o-Phenylenediamine dihydrochloride, Sigma-Aldrich, Merck KGaA, Darmstadt, Germany). The absorbance was measured at 450 nm using a Multiskan FC microplate reader (ThermoFisher Scientific, USA). The assay was performed twice with each sample in triplicate.

**Cell immunohistochemistry (IHC)**

To investigate the specific binding of nanobodies, we created a cell array where 20 cell lines were coated on a glass slide and stained with nanobodies. Cells were harvested using 3.4 mM of EDTA in

PBS, pelleted and resuspended in ddH2O. 7,500 cells were spotted per well on a 12-well black microscope cavity slides (Marienfeld, Germany). Cells were dried overnight in a fume hood. On the following day, slides were rehydrated in PBS 1x for 5 minutes, loaded on a Sequenza tray and blocked with 1 ml 1% BSA-PBS twice, each 30 min. Nanobody-phages were diluted in 150 ul 1% BSA-PBS to a final concentrations of 5E+10 PFU/slide and added twice, each 30 min. Slides were incubated with either mouse anti-PSMA monoclonal antibody (Sigma-Aldrich, USA) or NoNb phages as positive and negative controls, respectively. After 3 times washing with 0.2% BSA-PBS, slides were incubated with mouse monoclonal anti-M13 antibody (Abcam, USA). Binding of phages to the cells was detected by goat anti-mouse HRP antibody (Dako, Agilent, USA) followed by DAB substrate (Dako, Agilent, USA). Slides were visualized under light microscope and images from each well were captured using the camera (OLYMPUS, Japan) attached to the microscope. Staining intensity was scored on a scale from 0 to 4 (0: negative; 1: weakly positive; 2: medium positive; 3: highly positive; 4: strong positive). The scoring was conducted independently by two molecular biologists in a blinded manner. In cases of scoring discrepancies, the average score was used.

**Lentiviral transduction for PSMA knockdown**

To knockdown PSMA expression in LNCaP cell line, we used small hairpin RNA (shRNA) technology.

Five anti-PSMA shRNA plasmids (Mission shRNA libraries Sigma-Aldrich, Center for Biomics, Erasmus MC) were used in which the shRNA has been cloned into pLKO.1 vector carrying puromycin resistance gene as the selection marker. Initially, the bacterial samples carrying the plasmids were cultured on LB agar/ampicillin (100 ug/ml) plates. Single colonies were grown in LB/ampicillin (100 ug/ml) medium for plasmid DNA isolation using LabNed plasmid Midiprep kit (ThermoFisher Scientific Baltics UAB, Vilnius, Lithuania) following manufacturer's instructions.

For lentiviral production, HEK293T cells were transfected with a combination of 5 different shRNAs (total 20 ug) together with 15 ug packaging plasmid pPAX2 and 10 ug envelope plasmid pMD2.G using calcium phosphate transfection kit (Invitrogen, ThermoFisher Scientific, USA). Cells transfected with Lentivector carrying a CMV-driven Luc2-eGFP fusion (R980-M21-685) [6] and a non-targeting human gene shRNA (shNC) served as positive and negative control, respectively. Transfection efficiency was monitored using the GFP signal in Luc2-eGFP-transfected cells using fluorescence microscope (Nikon Ts2, China). The supernatant containing viruses was collected at 48 hrs post-transfection, centrifuged, filtered through a MF-Millipore Membrane filter 0.45 uM (Merck, US) and used to infect LNCaP cells. 48-72 hrs post infection, LNCaP cells went under puromycin selection with a concentration of 2.5 µg/mL for at least one week to ensure stable integration of the shRNA constructs.

**Western blot analysis of PSMA expression**

Cells were lysed using RIPA buffer (50mM Tris-HCl, 150 mM NaCl, 1% Triton X-100, 0.25% Sodium deoxycholate, 0.1% SDS, 2 mM EDTA) supplemented with phosphatase and protease inhibitors Complete (20X, Roche, Switzerland). The lysates were centrifuged at 14,000 rpm for 20 minutes at 4°C to remove debris, and supernatants were collected. Protein concentrations were determined using Pierce BCA protein assay kit (ThermoFisher Scientific, USA). 5 ug of each protein sample including wild type (WT), knockdown (shPSMA) and shRNA negative control (shNC) LNCaP cells along with DU145, B16-WT and B16-PSMA were resolved on a 10% SDS-PAGE gel and then transferred to a PVDF membrane (Millipore Sigma, Germany) using a semi-dry transfer apparatus (Bio-Rad, USA). Membranes were blocked with 5% skim milk in Tris-buffered saline with 0.1% Tween-20 (TBS-T) for 1 hour at room temperature. Subsequently, membranes were incubated overnight at 4°C with either rabbit monoclonal anti-PSMA antibody (Abcam, USA) or anti-β-actin antibody (ThermoFisher Scientific,

USA) and anti-human GAPDH antibody (Santa Cruz Biotechnology, USA) as controls. Protein bands were detected using goat-anti-rabbit HRP or goat-anti-mouse HRP (Dako, Agilent, USA) antibody for 1 hour at room temperature followed by incubating with BM chemiluminescence blotting substrate (Roche, Switzerland) and imaged using ChemiDoc Imaging System (Bio-Rad Laboratories, USA). Each experiment was repeated three times independently to ensure reproducibility.

**Binding of nanobodies to LNCaP cells in a flow cytometry assay**

Binding of nanobody-phages to wild type (WT) and knockdown (shPSMA) LNCaP cells were investigated in a flow cytometry assay. A7 and PSMANb9 nanobody-phages were directly labeled with 0.25 mM ATTO488 NHS-ester dye (ATTO-TEC, Germany) in 0.1 M NaHCO3, pH 8 for 3 hrs at room temperature. WT and shPSMA LNCaP cells were harvested using 3.4 mM EDTA, pelleted and resuspended in FACS buffer (10% FCS/PBS). 5E+10 labeled nanobody-phages were incubated with 500,000 cells on ice for 1 hour. LNCaP-WT cells stained with mouse monoclonal anti-PSMA antibody followed by goat-anti-mouse IgG cross-adsorbed secondary antibody Alexa Fluor 647 (AF647; Thermofisher Scientific, USA) and stained with secondary antibody only served as positive and negative controls, respectively. Unstained WT cells were used for gating. ATTO488 and AF647 signals were detected on a BD FACSSymphony A1 flow cytometer (BD Biosciences, USA) using AF488 and APC corresponding laser lines and filters, respectively. Raw data was analyzed using FlowJo software (version 10, BD Biosciences, US).

**Nanobody protein production**

A7 and PSMANb9 nanobodies were cloned into the pHEN vector (Twist Bioscience, US) and proteins expressed including a C-terminal Myc-tag and 6xHis tag using *E. coli* HB2151 cells as host. For protein expression, a single colony was cultured overnight and then diluted 1:100 in 2 liters fresh LB medium with 100 ug/ml ampicillin and cultured at 37°C until the OD600 reached 0.6-0.8. Isopropyl β-D-1-thiogalactopyranoside (IPTG) (Sigma-Aldrich, Germany) was added at a final concentration of 1 mM and the culture was incubated at 28°C overnight with 270 rpm shaking. Cells were harvested by centrifugation at 4,000xg for 30 minutes at 4°C. The cell pellets were resuspended in lysis buffer (50 mM $KH_2PO4$, 400 mM NaCl, 100 mM KCl, 10% glycerol, 0.5% Triton X-100, 8 M Urea, 1xProtease inhibitor, pH 8.0) and subjected to sonication (5x 20 seconds with 40 seconds off-time at 70% amplitude) on ice to lyse the cells. The lysate was cleared at 15,000xg for 30 minutes at 4°C and the supernatant was filtered through a 0.45 µm filter (Merck, US) to further clarify the lysate.

The lysate was then loaded onto a Ni-NTA agarose column (5 ml, Cytiva, USA) pre-equilibrated with wash buffer (50 mM $KH_2PO4$, 400 mM NaCl, PH 8.0) containing 25 mM imidazole. The column was washed with wash buffer with 25-100 mM imidazole to remove non-specifically bound proteins. The nanobodies were eluted using elution buffer (50 mM NaH2PO4, 300 mM NaCl, 250-500 mM imidazole, pH 8.0). The eluted nanobodies were concentrated and buffer exchanged with PBS using 10 KDa vivaspin6 MWCO column (2-6 mL, Cytiva, US) and the purity was assessed using mPAGE® 8–16% Bis-Tris Precast Gels (Merck KGaA, Darmstadt, Germany) followed by Coomassie Brilliant Blue staining. The protein concentration was then measured using Pierce BCA protein assay kit (Thermo Scientific, US), with bovine serum albumin (BSA) as a standard. Nanobody proteins were aliquoted in 50% (V/V) of glycerol and stored at -20°C.

***In silico* modeling of nanobody 3D structure**

The 3D structure of human prostate specific membrane antigen (PSMA) ectodomain (PDB ID: 1Z8L) was obtained from RCSB Protein Data Bank (https://www.rcsb.org/). The structure was processed to remove water molecules and unbound atoms using BIOVIA Discovery Studio Visualizer (Biovia, Dassault Systems, San Diego, CA). The 3D structures of five candidate nanobodies- A3, A7, PSMANb5, PSMANb6, PSMANb9, PSMANb11, were predicted using Robetta server (http://robetta.bakerlab.org) RoseTTAFold option that uses deep learning method, combines the advantages of template-based modeling and ab initio modeling and the SWISS-MODEL server (https://swissmodel.expasy.org) which use a homology modeling method [7]. The predicted models were downloaded and evaluated for their overall quality using multiple structure validation tools including ERRAT (https://saves.mbi.ucla.edu/) and PROSA-web (https://prosa.services.came.sbg.ac.at) which analyze non-bonded atomic interactions or local residue energy, respectively, compared to high resolution crystallographic structures in PDB. Additionally, the steric quality was assessed using Molprobity (http://molprobity.biochem.duke.edu) and PROCHECK (https://saves.mbi.ucla.edu/). Molprobity combines clash score, Rotamer outliers and Ramachndran analysis to validate the structure quality reporting a MolProbity score where good models typically have scores below 2.0. PROCHECK takes into account chain parameters and non-bonded interaction in addition to Ramachandran analysis, providing a quantitative table of percentage of residues in favored/allowed/disallowed Ramachandran regions and overall geometry deviations. ERRAT assesses the overall quality of protein models, reporting an ERRAT score (%) where scores within the range of 80-90% are considered as acceptable structure.

**Molecular docking**

The selected nanobody models and PSMA ectodomain were prepared for docking simulation using Dock Prep tool in UCSF Chimera (Resource for Biocomputing, Visualization, and Informatics (RBVI), University of California, San Francisco, USA), by adding hydrogens and assigning atomic charges. Key residues in the Complementarity Determining Regions (CDRs) were identified as potential binding sites. Nanobodies were docked to a monomer form of PSMA using the web “easy interface” of HADDOCK 2.4 (High Ambiguity Driven protein-protein DOCKing) web server (https://wenmr.science.uu.nl/haddock2.4). HADDOCK2.4 is a flexible docking software that integrates experimental biochemical or biophysical data to guide the docking of biomolecules [8] and has shown strong performance in multiple CAPRI (Critical Assessment of PRedicted Interactions) rounds [9, 10]. The generated models were clustered and ranked based on their HADDOCK score. To compare the PSMA epitopes bound by PSMA-617 and our selected nanobodies, we used PyRx 0.8 [11] and Autodock 4.2 (The Scripps Research Institute, La Jolla, CA, USA) [12] to dock PSMA617 peptide to PSMA ectodomain. Molecular interactions were visualized and analyzed using Python Molecular Graphics (PyMOL, Schrödinger, LLC, USA) and LigPlot+ software [13].

**Molecular dynamics simulation**

Molecular dynamics (MD) simulations were performed for 200 ns using GROMACS 2022 package (GROningen MAchine for Chemical Simulations, the Netherlands) and OPLS-AA force field (Optimized Potential for Liquid Simulations-All Atom, Yale University, USA) to evaluate the stability of the A7-PSMA and Nb9-PSMA complexes. To set up the system, the selected complexes were dissolved in a TIP3P water box and neutralized with $Na^+/Cl^-$ ions to address the charge imbalance caused by ligand-receptor interactions and solvation. Notably, ligand-receptor interactions may alter the total charge, and the addition of water molecules can shift the system's charge from zero. During simulation, the system was neutralized by adding $Cl^-$ ions if the net charge was positive or $Na^+$ ions if it was negative. Energy minimization was performed, followed by equilibration in two stages: first, under the NVT ensemble at 300 K for 100 ps, and then under the NPT ensemble at 300 K and 0.1 bar pressure for 100 ps using a Parrinello-Rahman barostat. Simulation parameters included long-range electrostatics calculated via the Particle Mesh Ewald (PME) algorithm with a 10 Å cutoff, van der Waals interactions with a 1 nm cutoff, and covalent bonds constrained using the LINCS algorithm. After equilibration, a 200 ns production MD simulation was conducted. The stability of the complexes was assessed by calculating the root mean square deviation (RMSD), root mean square fluctuation (RMSF), and radius of gyration (Rg).

**Flow cytometry**

To validate the *in silico* predictions, a titration assay was conducted to determine the nanobody concentration sufficient to fully block the PSMA binding sites on the cell surface. 500,000 LNCaP or B16-PSMA cells were incubated with increasing concentrations (125, 250, 500, 1000, and 2000 µg/mL) of A7 or PSMANb9 nanobody proteins for 1 hour in 5% FCS-PBS on ice. Binding of nanobody protein was then detected using a mouse monoclonal anti-myc-tag antibody (Sigma-Aldrich, Germany) followed by goat-anti-mouse IgG cross-adsorbed secondary antibody Alexa Fluor 647 (Thermofisher Scientific, USA). Similar titration assay was performed with varying concentrations (1E+11, 5E+10, 1E+10, 5E+9, 1E+9, and 5E+8) of ATTO488-labeled A7 or PSMANb9 nanobody-phages. For competition assay, LNCaP or B16-PSMA cells were initially blocked with an appropriate concentration of A7 or PSMANb9 proteins followed by incubation with optimized concentration of ATTO488-labeled A7 or PSMANb9 nanobody-phages. As control, cells were incubated with either ATTO488-labeled A7 or PSMANb9 nanobody-phages, alone. To investigate the competition between A7 and PSMANb9 proteins, cells were first incubated with A7 or PSMANb9 proteins followed by adding ATTO488-labeled PSMANb9 or A7 nanobody-phages, respectively. No washing was performed between the two incubations. Signal detection and data analysis was performed following the protocol described above.

#### References


1. Chatalic, K.L., et al., *A Novel $^{111}$In-Labeled Anti-Prostate-Specific Membrane Antigen Nanobody for Targeted SPECT/CT Imaging of Prostate Cancer.* J Nucl Med, 2015. **56**(7): p. 1094-9.
2. Ghandi, M., et al., *Next-generation characterization of the cancer cell line encyclopedia.* Nature, 2019. **569**(7757): p. 503-508.
3. De Morrée, E.S., et al., *Loss of SLCO1B3 drives taxane resistance in prostate cancer.* British journal of cancer, 2016. **115**(6): p. 674-681.
4. Li, M., et al., *Sequence and insertion sites of murine melanoma-associated retrovirus.* Journal of virology, 1999. **73**(11): p. 9178-9186.
5. Fadrosh, D.W., et al., *An improved dual-indexing approach for multiplexed 16S rRNA gene sequencing on the Illumina MiSeq platform.* Microbiome, 2014. **2**(1): p. 6.
6. Day, C.P., et al., *Lentivirus-mediated bifunctional cell labeling for in vivo melanoma study.* Pigment cell & melanoma research, 2009. **22**(3): p. 283-295.
7. Baek, M., et al., *Accurate prediction of protein structures and interactions using a three-track neural network.* Science, 2021. **373**(6557): p. 871-876.
8. Dominguez, C., R. Boelens, and A.M.J.J. Bonvin, *HADDOCK: a protein− protein docking approach based on biochemical or biophysical information.* Journal of the American Chemical Society, 2003. **125**(7): p. 1731-1737.

9. Reys, V., et al., *Integrative Modeling in the Age of Machine Learning: A Summary of HADDOCK Strategies in CAPRI Rounds 47–55.* Proteins: Structure, Function, and Bioinformatics, 2024.
10. Koukos, P.I., et al., *An overview of data-driven HADDOCK strategies in CAPRI rounds 38-45.* Proteins: Structure, Function, and Bioinformatics, 2020. **88**(8): p. 1029-1036.
11. Dallakyan, S. and A.J. Olson, *Small-molecule library screening by docking with PyRx*, in *Chemical biology: methods and protocols*. 2014, Springer. p. 243-250.
12. Morris, G.M., et al., *AutoDock4 and AutoDockTools4: Automated docking with selective receptor flexibility.* Journal of computational chemistry, 2009. **30**(16): p. 2785-2791.
13. Wallace, A.C., R.A. Laskowski, and J.M. Thornton, *LIGPLOT: a program to generate schematic diagrams of protein-ligand interactions.* Protein engineering, design and selection, 1995. **8**(2): p. 127-134.
